# ATRX alteration contributes to tumor growth and immune escape in pleomorphic sarcomas

**DOI:** 10.1101/2020.10.23.352112

**Authors:** Lucie Darmusey, Gaëlle Pérot, Noémie Thébault, Sophie Le Guellec, Nelly Desplat, Laëtitia Gaston, Lucile Delespaul, Tom Lesluyes, Elodie Darbo, Anne Gomez-Brouchet, Elodie Richard, Jessica Baud, Laura Leroy, Jean-Michel Coindre, Jean-Yves Blay, Frédéric Chibon

**Author notes:** Correspondence: Frédéric Chibon, Cancer Research Center in Toulouse (CRCT), 2 avenue Hubert Curien, 31037, Toulouse, France, 0582741765. These authors contributed equally (Lucie Darmusey, Gaëlle Pérot).

## Abstract

Whole genome and transcriptome sequencing of a cohort of 67 leiomyosarcomas revealed *ATRX* to be one of the most frequently mutated genes in leiomyosarcomas after *TP53* and *RB1*. While its function is well described in the alternative lengthening of telomeres mechanism, we wondered whether its alteration could have complementary effects on sarcoma oncogenesis. *ATRX* alteration is associated with the down-expression of genes linked to differentiation in leiomyosarcomas, and to immunity in an additional cohort of 60 poorly differentiated sarcomas. *In vitro* and *in vivo* models showed that *ATRX* loss increases tumor growth rate and immune escape by decreasing the immunity load of active mast cells in sarcoma tumors. These data indicate that an alternative to unsuccessful targeting of the adaptive immune system in sarcoma could be to target the innate system. This might lead to a better outcome for sarcoma patients in terms of *ATRX* status.

## Introduction

Pleomorphic sarcomas are a group of rare mesenchymal tumors comprising different histotypes such as undifferentiated pleomorphic sarcoma (UPS), myxofibrosarcoma (MFS), dedifferentiated liposarcoma (DDLPS), osteosarcoma (OS) and leiomyosarcoma (LMS), which is the most frequent subtype ^1^. LMS has a smooth muscle differentiation and can occur in any anatomical site, although there are three main locations: limbs, trunk and uterus. Currently, the first-line treatment is wide-margin resection for localized tumors and anthracycline-based chemotherapy for advanced tumors ^2^. However, these treatments are still not effective enough as 48 to 89% of LMS develop metastases depending on the tumor location and the mortality rate is between 50 and 65% with a median survival of around 12 months ^3,4^. From a genomic standpoint, LMS, like other pleomorphic sarcomas, have a very rearranged and unbalanced karyotype ^2^.

In a whole genome and whole transcriptome sequencing study conducted by our team, Darbo *et al.* showed that LMS could be separated into two groups with specific clinical, transcriptomic and genomic features: the homogenous and the other LMS. Those groups share a low somatic mutation burden and a high level of copy-number alterations ^5^. But only three genes came out to be recurrently mutated (considering point mutations only), as also showed by the TCGA study ^6^: *TP53*, *RB1* and *ATRX* (mutated in 48.7%, 17.9% and 12.8% respectively). *RB1* and *TP53* are tumor suppressor genes that have long been known to be implicated in the oncogenesis of pleomorphic sarcomas ^7–10^. *ATRX* is a chromatin modifier gene with a Swi/Snf2 domain ^11^. Its tumor suppressive function has so far been related to its role in the alternative lengthening of telomeres (ALT) mechanism ^12^, inducing genome instability ^13^ and leading to a poor prognosis of *ATRX*-altered tumors ^14^. Recently, its involvement in senescence ^15^ and in intrinsic immunity *via* its interactions within promyelocytic leukemia nuclear bodies (PML NBs) ^16^ was questioned. Here, we investigated whether ATRX might have additional impacts in the oncogenesis of pleomorphic sarcomas beyond its role in the ALT mechanism and show how its involvement in oncogenesis is also linked to differentiation, tumor growth and immunity.

## Results

### Distinct genetic alterations trigger loss of ATRX protein

Sixty-seven LMSs (Cohort 1; Table S1) were sequenced at the whole genome and transcriptome levels (28 LMSs + 39 LMSs from ^5^) and *ATRX* was identified as the third most frequently mutated gene after *TP53* and *RB1*. By integrating point mutations and structural variations (SV), ATRX is altered in 20 cases (29.8 %; Figure 1), with 8 point mutations (missense (MS) and non-sense (NS); 40%), 7 frameshifts (FS; 35%) and 5 structural variants (SV; 25%). All mutations and SV were validated by an independent technique (RNA sequencing and/or Sanger sequencing) (Tables S2 and S3). *ATRX* was altered in 23.7% of non-uterine LMS (14/59) compared to 75% of uterine LMS (6/8), which is significantly higher in this specific anatomical site (P = 7.002 × 10^−3^; Figure 2A and Figure S1). Altered cases were not enriched in any other clinical annotation (*i.e*. grade, metastasis or sex). Regarding SV, 3 out of 5 led to a loss of *ATRX* expression and the other two led to a frameshift (Table S2). These alterations were hemizygous in the three males due to the location of *ATRX* on chromosome X (Xq21), and in two females with either a deletion of the second allele (LMS69) or an isodisomy (LMS49). In the other 15 females, 93.3% of the alterations (14/15) occurred on the active X, as RNAseq analysis showed the altered transcript expression (Table S2). No expression of the mutated allele was detected in LMS48 (Figure 2A and Table S2). Alterations were distributed throughout the whole gene but two regions were most frequently affected: one between exons 17 and 21 (40%, 6/15) and the other in exon 9 (33.4%, 5/15). At the mRNA level, mutated cases had a significantly lower *ATRX* expression than wild-type (WT) tumors (P = 3.79 × 10^−4^; Figure 2B) and at the protein level, alterations led to a loss of nuclear protein (P = 8.04 × 10^−10^; Figure 2B and Figure S2).

**Figure 1:**
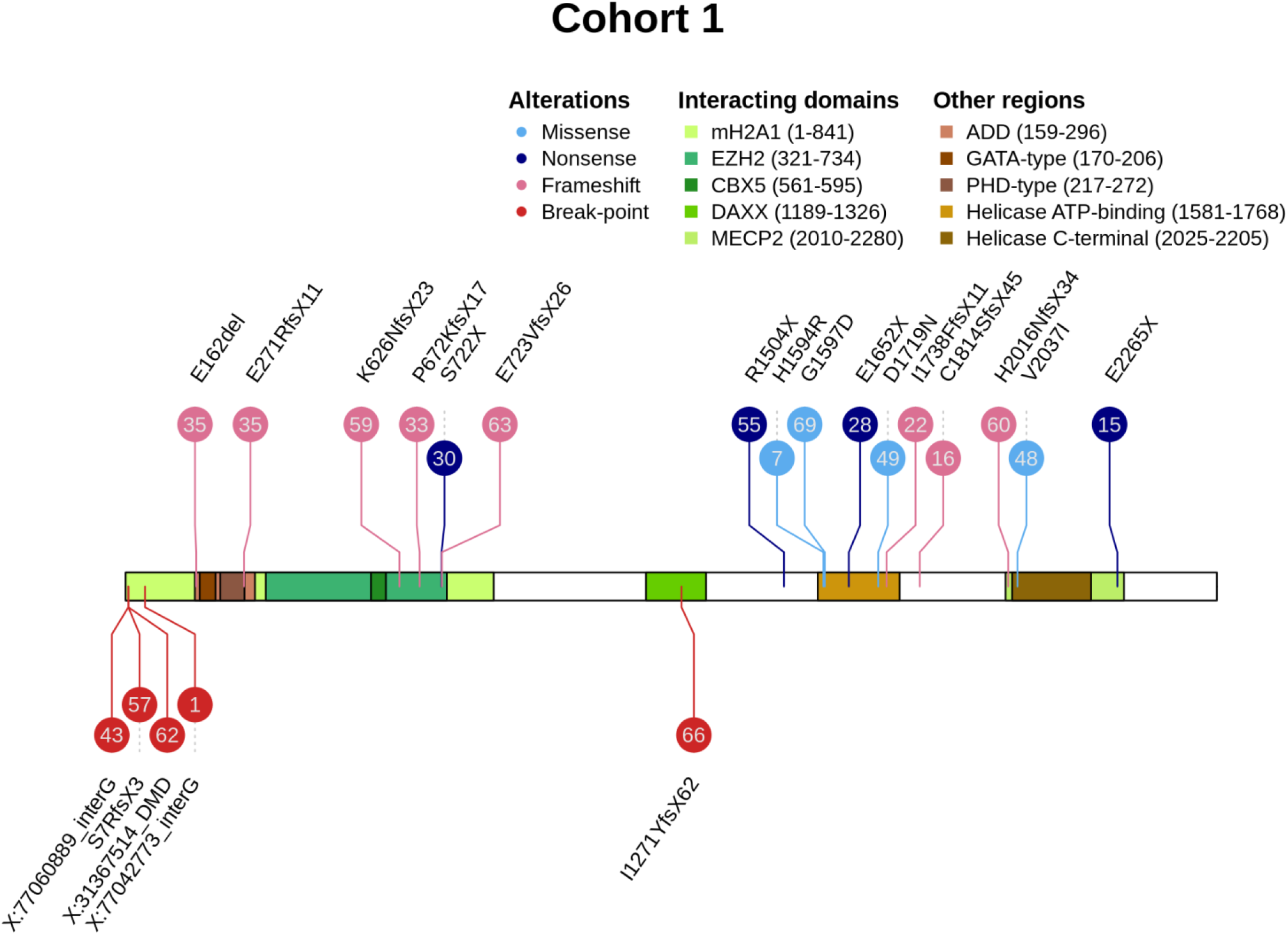
*ATRX* mutations and structural variants *ATRX* alterations are color-coded according to their type (legend at the top). Numbers in bubbles represent tumor sample. Consequences of all point mutations on ATRX protein are annotated above a schematic representation of the protein, or below for two structural variants. See also Table S1 and S3.

**Figure 2:**
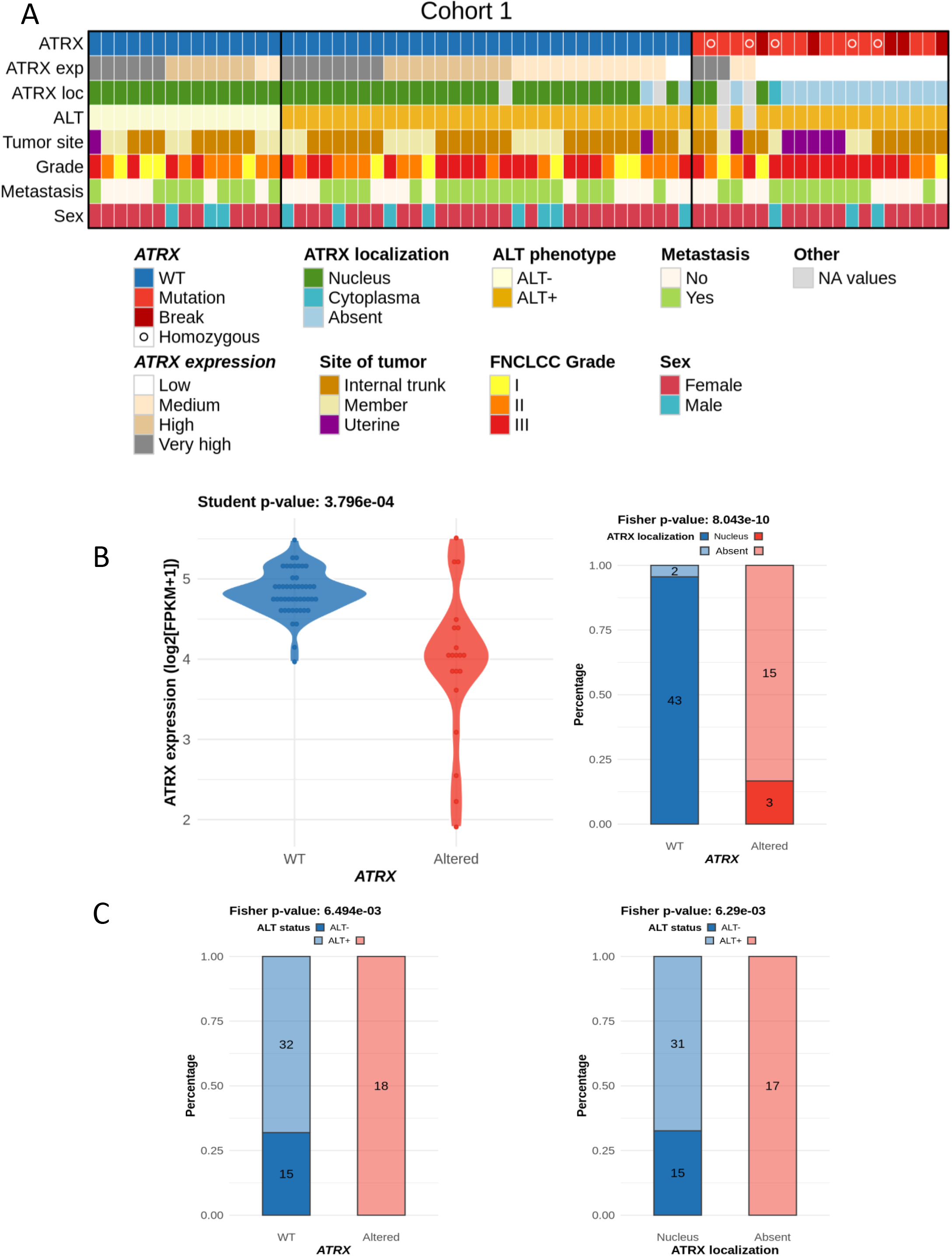
ATRX status and integrated representation **A** Integrated representation shows *ATRX* alterations, *ATRX* mRNA expression (by quartile), ATRX localization, ALT mechanism phenotype, tumor site, FNCLCC grade, presence or not of metastasis and sex of each patient. Tumors are ordered by *ATRX* status, ALT phenotype, mRNA expression and protein localization. **B** Association between *ATRX* alteration and its mRNA expression (log2(FPKM+1)) (left) or its protein localization (right). **C** Relation between *ATRX* status (left) or its protein localization (right) and ALT mechanism phenotype. For ATRX localization, the “absent” group means “not at the nucleus”, including all cases without expression and the case with a cytoplasmic localization (LMS16). P-values were calculated with Student test for **B - left** and with Fisher test for **B - right** and **C**. See also Table S1 and S2, Figure S1, S2, S3 and S4.

### ATRX alteration is linked to ALT mechanism

Since *ATRX* loss is linked to the ALT phenotype ^17^, the ALT status of tumors was determined. Most LMS were ALT-positive (ALT+, 76.9%, 50/65) (Figure 2A and Figure S3). Both *ATRX* alteration (P = 6.49 × 10^−3^) and ATRX protein loss (P = 6.29 × 10^−3^) were significantly associated with the ALT mechanism (Figure 2C). However, while all *ATRX*-altered cases were ALT+, most ALT+ cases were *ATRX* WT (64%, 32/50) with 93.3% of cases (28/30) expressing the protein in the nucleus (Figures 2A and 2C).

### *ATRX* alteration is not associated with prognosis

Neither *ATRX* status (altered or WT), mRNA expression (below or above defined cut-off, see material and methods section), protein localization (nuclear or absent), nor ALT phenotype (positive or negative) could split patients into two groups with significantly distinct prognoses (Figure S4).

### Differentiation transcriptomic programs is modified upon *ATRX* alteration

Searching for the oncogenic impact of these *ATRX* alterations, we tested whether altered tumors had a distinct transcriptomic program and identified 340 and 219 genes significantly down- and up-expressed in the *ATRX*-altered group, respectively (P < 0.05; Figure 3A). Functional enrichment analysis (Figure 3A) showed that genes down-expressed were significantly involved in blood pressure, heart contraction and striated muscle contraction. These findings were strengthened when patients were grouped according to ATRX protein localization, since genes down-expressed upon protein loss were found to be involved in similar biological mechanisms, *i.e*. muscle system and contraction (Figure 3B).

**Figure 3:**
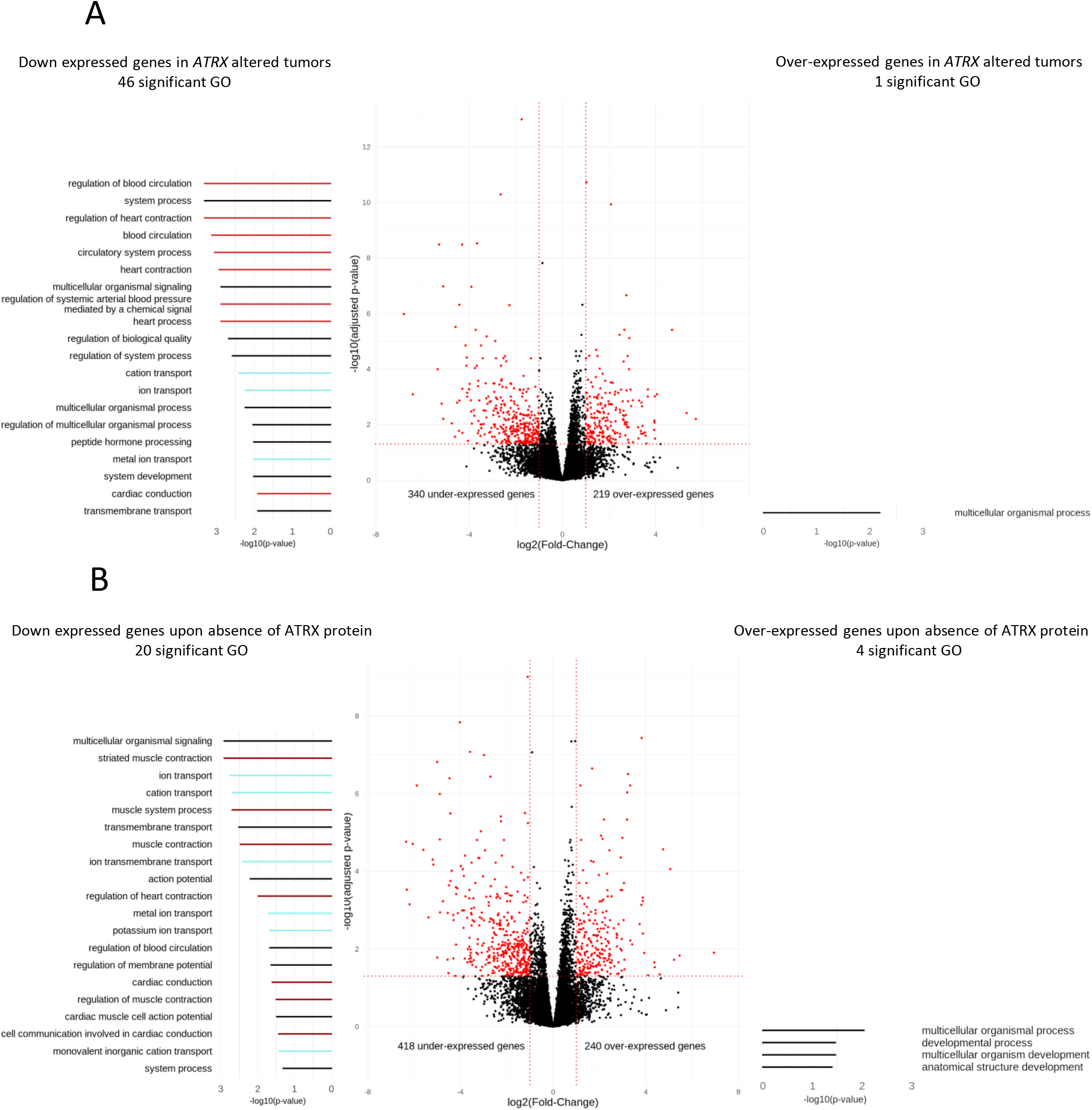
Differential gene expression and Gene Ontology analyses according to ATRX alteration in leiomyosarcomas (Cohort 1) Differentially expressed genes according to **A** *ATRX* status (wild-type *vs* altered) or **B** ATRX expression (nucleus *vs* absent). Red dots indicate significant genes (*P* ≤ 0.05 and fold-change ≤ −2 or ≥ 2). Gene Ontology (GO) analyses, represented on the left (under-expressed genes) and the right (over-expressed genes), identified 46 and 1 significant GO terms (*P* ≤ 0.05), respectively in **A** and 20 and 4 significant GO terms (*P* ≤ 0.05) in **B**. On each side, the 20 most significant GO terms are represented and color-coded by mechanisms; light red, dark red, light blue and black colors indicate “circulatory system process”, “muscle system process”, “ion transport” and general terms, respectively. For ATRX localization, the “absent” group means “not at the nucleus”, including all cases without expression and the case with a cytoplasmic localization (LMS16). All p-values adjusted by Benjamini and Hochberg method. See also Figure S5.

As expected, clustering based on these 559 differentially expressed genes (Figure S5A) revealed a group with a high percentage of *ATRX*-altered patients (75%, 15/20). Patients in this cluster had tumors that were enriched in uterine or “other” LMS type (P = 3.49 × 10^−7^; Figure S5B)^5^. “Other” LMS are less differentiated than “homogeneous” LMS and are thought to derive from fibroblasts rather than smooth muscle cells (SMC) ^5^.

The association between enrichment of down-expressed genes linked to muscle system and of oLMS in ATRX altered tumors suggested that either *ATRX* alteration preferentially occurs in partially or undifferentiated cells, or that it may induce dedifferentiation. To explore these hypotheses, we studied the *ATRX* status in a second cohort comprising poorly differentiated pleomorphic sarcomas characterized by RNAseq.

### *ATRX* alterations are recurrent and similar in poorly differentiated pleomorphic sarcomas

RNA sequencing of 60 pleomorphic sarcomas (cohort 2; Table S1) from a previously published cohort ^18^ was reanalyzed and 10 *ATRX*-altered tumors (16.7%) were identified (Tables S2 and S4). The types of alteration as well as their functional consequences were similar to those detected in cohort 1 (Figure 4A). Altered cases were not enriched in any annotation (*i.e*. histotype, tumor site, grade, metastasis or sex) (Figure 4B) but had a significantly lower mRNA expression of *ATRX* (P = 3.62 × 10^−2^; Figure 4C) and were significantly associated with ALT (P = 3.96 × 10^−3^; Figure 4D). *ATRX*-altered tumors did not have a distinct prognosis in cohort 2 (Figure S6A), nor when the two cohorts are merged (Figure S6B).

**Figure 4:**
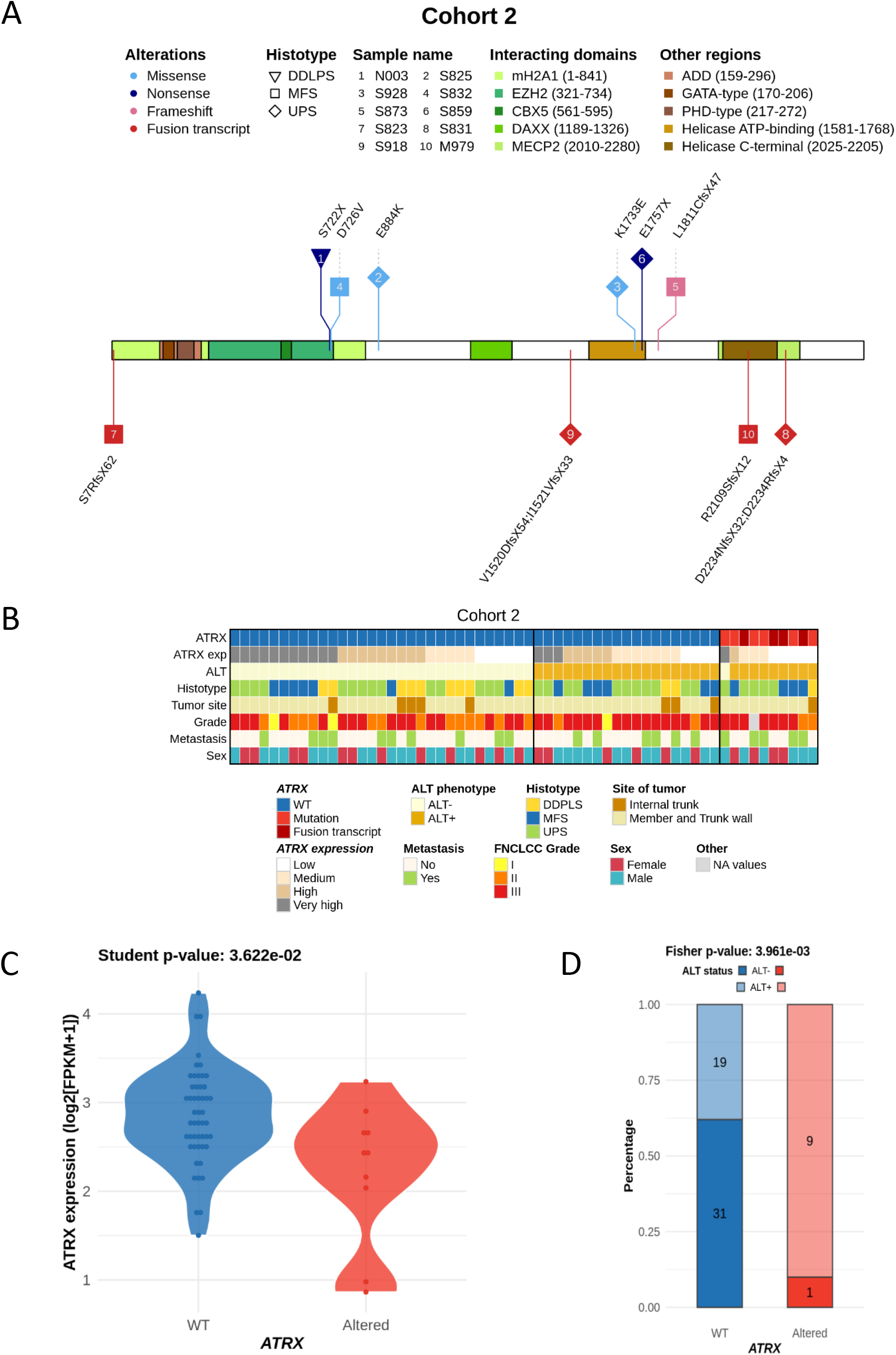
*ATRX* alterations and integrated representation in poorly differentiated pleomorphic sarcomas (cohort 2) **A** *ATRX* alterations are color-coded by their type and shapes represent histotypes. Numbers in bubbles indicate a tumor sample (legend at the top). Translated consequences on ATRX protein is annotated above a protein schematic representation for mutations, or below for fusion transcripts. **B** Integrated representation shows *ATRX* alterations, mRNA expression (by quartile), ALT mechanism phenotype, histotypes, tumor site, FNCLCC grade, presence or not of metastasis and sex of each patient. Tumors are ordered by *ATRX* status, ALT phenotype, mRNA expression and histotypes. **C** Association between *ATRX* status and its mRNA expression (log2(FPKM+1)). **D** Relation between *ATRX* status and ALT phenotype. See also Table S1, S2, S4 and Figure S6.

### Immunity transcriptomic program is modified upon *ATRX* alteration in poorly differentiated sarcomas

Functional enrichment analysis of differentially expressed genes showed that *ATRX* alteration induced the overexpression of 76 genes enriched in GO terms related to the metabolic process, and the down-expression of 506 genes enriched in GO related to immunity. The five most significantly enriched GO were (Figure 5) “T cell activation” (P = 8.41 × 10^−19^), “lymphocyte activation” (P = 1.13 × 10^−17^), “immune system process” (P = 4.82 × 10^−16^), “leukocyte activation” (P = 1.58 × 10^−14^) and “regulation of immune system process” (P = 1.82 × 10^−14^).

**Figure 5:**
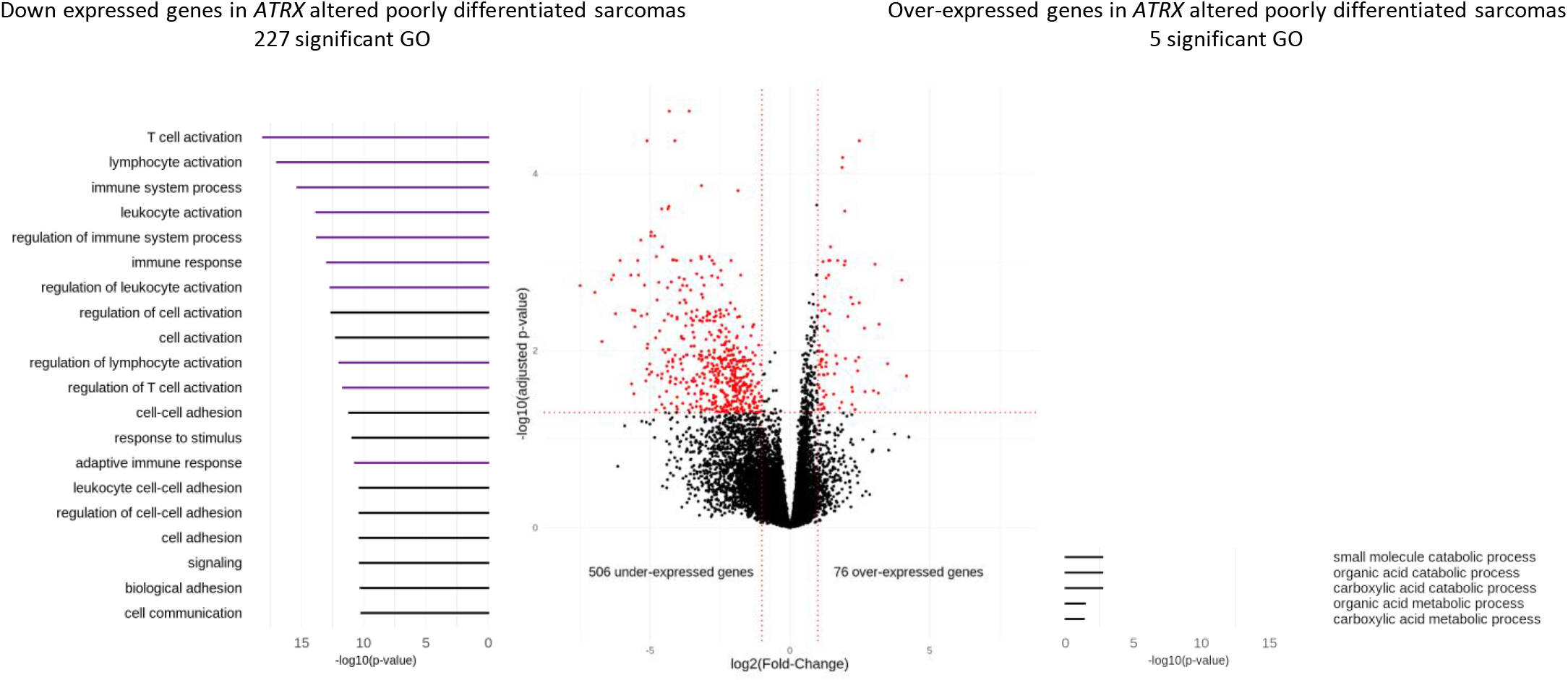
Differential gene expression and Gene Ontology analyses according to *ATRX* status (wild-type *vs* altered) in poorly differentiated sarcomas (Cohort 2) Differentially expressed genes in *ATRX*-altered tumors are represented in red (*P* ≤ 0.05 and fold-change ≤ −2 or ≥ 2). Gene Ontology (GO) analyses, represented on the left (under-expressed genes) and the right (over-expressed genes), identified 227 and 5 significant GO terms (*P* ≤ 0.05), respectively. On the left, the 20 most significant GO terms are represented and color-coded by mechanism; purple and black groups indicate “immunity system process” and general terms, respectively. All p-values adjusted by Benjamini and Hochberg method.

Results from both cohorts indicated that *ATRX* alteration is associated with differentiation and immunity. Since this is particularly relevant as immunotherapies are currently not efficient in sarcomas, we functionally tested the hypothesis that *ATRX* alteration might modify the anti-tumor immune response.

### *ATRX* knock-down impact oncogenic features toward aggressiveness

To functionally test the impact of *ATRX* alterations, three models of *ATRX* knock-down (*ATRX^KD^*) were constructed: i) a model to evaluate tumor growth *in vitro* and *in vivo* in a human UPS cell line (IB106), ii) another to study immunity in immunocompetent mice (Balb/c) with a mouse poorly differentiated OS (K7M2; a sarcoma with very close genetics to UPS and LMS) and iii) a third to compare human and mouse, using a human OS cell line (MG63). These cell lines were transducted by lentivirus with an *ATRX* shRNA. Western blot evidenced the successful extinction of ATRX in each cell line (Figure 6A). ALT analysis showed that *ATRX* shRNA did not change ALT status in any cell line (Figure S7A).

**Figure 6:**
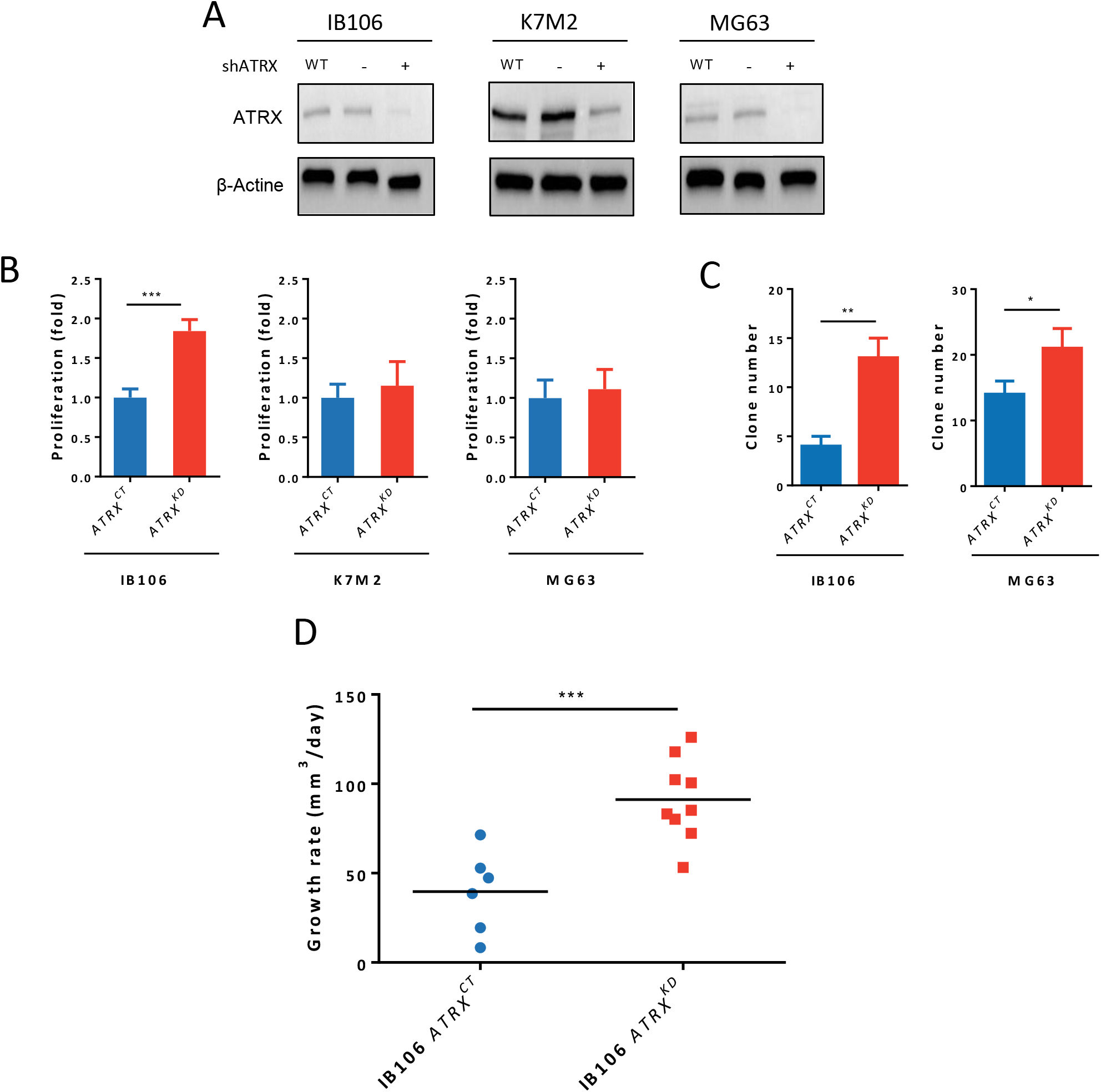
*ATRX* knock-down increases aggressiveness of sarcoma cells **A** *ATRX* knock-down by shRNA validation in western blot in K7M2, MG63 and IB106 cell lines. **B** Proliferation analysis by MTT after 4 days, comparing *ATRX^CT^* and *ATRX^KD^* cells in K7M2, MG63 and IB106 cell lines (Data are represented as mean ± s.d.; n = 3 independent experiments). **C** Soft agar assay analysis comparing *ATRX^CT^* and *ATRX^KD^* cells in K7M2, MG63 and IB106 cell lines (Data are represented as mean ± s.d.; n = 4 independent experiments). Images were taken after 4 weeks and crystal violet staining. **D** Tumor growth rate analysis of IB106 *ATRX^CT^* or IB106 *ATRX^KD^* cells sub-cutaneous xenografts on NSG mice (N = 10 in each group). Growth rate is calculated by segmental linear regression with GraphPad. The black line represent the mean. \**P* ≤ 0.05, \*\**P* ≤ 0.01, \*\*\**P* ≤ 0.001, p-value was calculated with 2-way ANOVA for A and unpaired t-test for B, C and D. See also Figure S7A.

*In vitro*, a significant (P < 0.0001) increase in proliferation was observed in the UPS cell line IB106 *ATRX^KD^* but not in OS cell lines (K7M2 and MG63) (Figure 6B). Colonies formed in soft agar assay revealed that the mouse cell line K7M2 was unable to form any colony with or without ATRX expression. In contrast, there was a significant increase in colony number in human cell lines IB106 and MG63 upon *ATRX* down-expression, from a mean of 14 to 21 colonies (P = 2.6 × 10^−3^) and from 4 to 13 (P = 1 × 10^−3^), respectively (Figure 6C). Next, IB106 *ATRX^CT^* (control) and *ATRX^KD^* cells were subcutaneously grafted in 10 NSG mice each. A tumor grew in 6/9 *ATRX^CT^* group and in 9/10 *ATRX^KD^* group. Tumor growth rates were three-fold higher in *ATRX^KD^* tumors (91.2 ±7.6 mm^3^/day) than in *ATRX^CT^* tumors (32.9 ±10.6 mm^3^/day) (P = 5 × 10^−4^, Figure 6D).

### ATRX knock down modifies anti-tumor immune response in vivo

The involvement of ATRX alteration in immune escape was tested by grafting K7M2 *ATRX^CT^* and *ATRX^KD^* cells in immunodeficient NSG mice and in immunocompetent Balb/c mice (N=15 for each group). Growth rate was not significantly increased upon ATRX knock-down in any hosts (Figure 7A). Tumor-free survival in *ATRX^CT^* and in *ATRX^KD^* models displayed no significant differences in immunodeficient NSG mice, whereas in immunocompetent Balb/c mice there was 53.4% (8/13) of tumor induction with K7M2 *ATRX^CT^ versus* 92.8% (13/14) with K7M2 *ATRX^KD^*. Therefore, tumor-free survival was significantly poorer upon *ATRX* knock-down (P = 9.7 × 10^−3^; Figure 7B).

**Figure 7:**
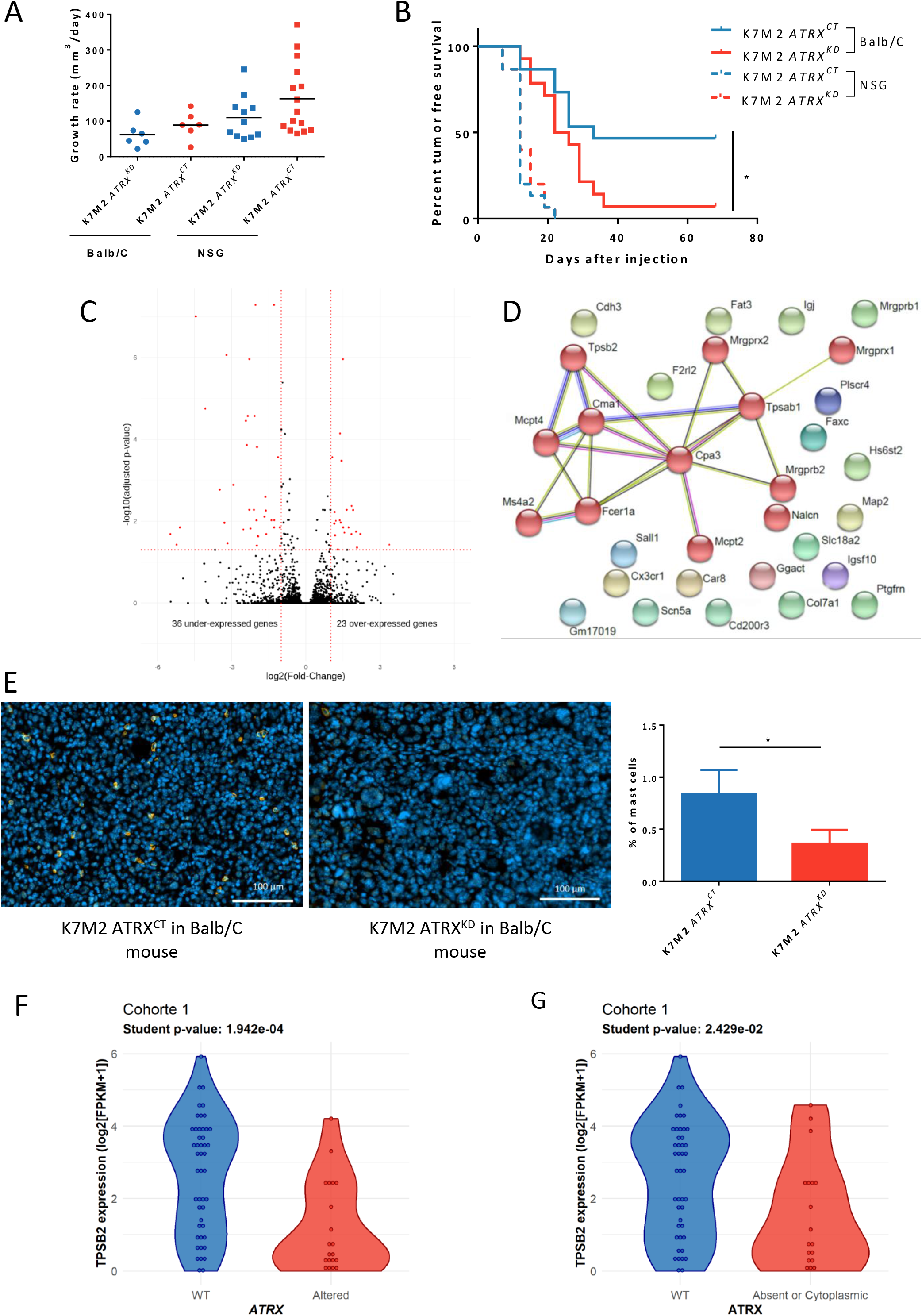
*ATRX* knock-down allows immune escape of sarcomas *via* non-recruitment of mast cells **A** Tumor growth rate analysis of K7M2 *ATRX^CT^* or K7M2 *ATRX^KD^* cells xenografted under the skin of NSG or Balb/c mice (N = 15 in each group). The black line represent the mean. **B** Tumor-free survival curves of K7M2 *ATRX^CT^* or *ATRX^KD^* tumors in immunodeficient NSG mice and immunocompetent Balb/c mice (N = 15 mice for each condition) using Kaplan-Meier method. **C** Comparison of RNA expression in log2(FPKM+1) of K7M2 *ATRX^KD^* tumors *versus* K7M2 *ATRX^CT^* tumors developed in immunocompetent mice (n =4 tumors in each condition) showing 23 and 37 significantly up- and down-expressed genes in K7M2 *ATRX^KD^* tumors, respectively. **D** Links between down-expressed genes in K7M2 *ATRX^KD^* tumors found by the STRING Database showing one cluster with genes involved in mast cells *via* MCL clustering. **E** Immunostaining of mast cells by targeting tryptase in K7M2 *ATRX^CT^* and K7M2 *ATRX^KD^* tumor tissues with nucleus marked with DAPI. On the right, percent of mast cells in the two conditions (Data are represented as mean ± s.d.; n = 4 tumors). **F** TPSB2 mRNA expression in log2(FPKM+1) according to *ATRX* status in cohort 1. **G** TPSB2 mRNA expression in log2(FPKM+1) according to ATRX localization in cohort 1. \**P* ≤ 0.05, \*\**P* ≤ 0.01, \*\*\**P* ≤ 0.001, p-value was calculated with Mantel-Cox test for B and unpaired t-test for E, F, G. See also Figure S7B.

Differential gene expression analysis between *ATRX^KD^ vs ATRX^CT^* K7M2 tumors in Balb/c mice revealed that 37 genes were down-regulated and 23 genes were overexpressed in *ATRX^KD^* tumors (Figure 7C). The low number of genes precluded any functional enrichment analysis. Consequently, a String Protein Interaction ^19^ analysis was performed. Whereas no consistent clusters arose with up-regulated genes upon *ATRX* knock-down (Figure S7B), one emerged in down-regulated genes, with 12 proteins out of 37 linked to mast cell pathways (including *TPSB2* coding tryptase, a widely used mast cells marker) (Figure 7D).

Immunofluorescence against tryptase on the murine tumors previously processed in RNAseq showed that mast cells expressing tryptase represented a mean of 0.8% of total cells in *ATRX*^CT^ tumors, whereas they constituted 0.3% of *ATRX*^KD^ tumors (P = 0.01; Figure 7E). This significant difference prompted us to assess whether the proportion of infiltrating mast cells in human sarcomas is also related to *ATRX* alteration and the absence of ATRX from the nucleus. In the LMS cohort, the only one fully characterized at every level, *TPSB2* was significantly under-expressed in *ATRX*-altered cases (P = 1.9 × 10^−4^; Figure 7F) and in tumors with no nuclear ATRX (P = 0.02; Figure 7G), which might indicate that there are fewer infiltrating mast cells in these human LMS.

## Discussion

This in-depth *ATRX* genetic analysis revealed that *ATRX* alteration likely affects a quarter of pleomorphic sarcomas, since it was found in 29.8% of LMS and in 16.7% of undifferentiated sarcomas (US). Cohort 2 is less deeply characterized (WGseq for cohort 1, RNAseq for cohort 2), so cases might be missed with this RNAseq-based screening (20.7% observed for UPS/MFS/DDLPS in TCGA ^6^). The rate of alteration in LMS is consistent with the rate of 24% found by Chudasama and colleagues ^20^, but it is slightly higher than that generally observed in other LMS cohorts, which is around 16% ^6,21,22^, probably due to the exhaustiveness of WGseq. *ATRX* mutations were distributed across the entire gene, as previously observed ^6,20,23,24^. Three main factors link the two types of sarcomas in the present study: i) in females, all alterations except in LMS48 can be interpreted as occurring on the active X, ii) point mutations are more frequent (75% in LMS and 60% in US) than structural variations (25% in LMS and 40% in US), as previously observed ^6,20^; and iii) the alterations lead preferentially to a frameshift and thus to a truncated protein in 66.7% of cases (20/30, 65% in LMS and 70% in US), in agreement with previous descriptions in sarcomas ^6,20,23,25^. Of note, *ATRX* alterations in the present study were not significantly associated with a poorer prognosis. However, this association depends on the cohorts studied ^14,24^ and was observed in only one cohort that mainly included missense mutations^24^.

*ATRX* mutated cases were also linked to the location of LMS, *i.e*. 75% of uterine cases were *ATRX*-altered (6/8). Loss of *ATRX* in uterine tumors is a key difference between benign and malignant tumors. In this location, it has been proposed to use *ATRX* loss as a marker of the highly probable evolution of benign tumors toward malignancy ^26^. In other LMS locations, ATRX loss is linked to the “other” LMS group. LMS belonging to this subtype are mainly poorly differentiated and likely originate from fibroblastic cells ^5^. Furthermore, as ATRX loss in LMS is associated with a lower expression of genes related to smooth muscle activity, we hypothesize that it occurs preferentially in poorly differentiated cells. The degree of cell differentiation may be crucial for the loss of ATRX to confer advantages to the precursor sarcoma cell.

*ATRX* knock-down modifies tumor cell proliferation, as confirmed *in vivo* where *ATRX* knock-down tumors grew three-fold faster than controls, and clonogenicity in sarcoma models. Interestingly, poorly differentiated sarcomas with *ATRX* alteration overexpressed genes related to metabolism, whose upregulation is a known hallmark of cancers and supports cell survival and proliferation ^27^. The hypothesis that ATRX could act through metabolism regulation is a very appealing one that now requires functional validation.

*In vivo* experimentation revealed a new role of ATRX, as its alteration was associated with a poorer outcome exclusively in an immunocompetent murine host, and with down-expression of immune-related genes in poorly differentiated human sarcomas. These two findings show that ATRX loss can influence the regulation of immune response in sarcomas, probably by limiting mast cell recruitment, as evidenced by the lower proportion of tumor infiltrating mast cells upon *ATRX* down-expression. The role of mast cells in tumor control is currently considered as dual and antagonistic, since they can support tumorigenesis or suppress tumor growth. Their role is dependent on the type of tumor ^28^. To our knowledge, no study has yet investigated the role of mast cells in the oncogenesis of sarcomas. *FcεRI* and *Ms4a2* are two down-expressed genes in *ATRX^KD^* K7M2 tumors. They are part of the IgE activating mast cell pathway that confers them a protective role in epithelial tumors ^29^. In addition to their higher proportion, these mast cells present in *ATRX^WT^* tumors likely play a suppressor role in which they recruit other immune cells to tumor sites by enhancing vascular permeability and direct chemoattraction ^30^. In human LMS, the absence of ATRX is linked to the down-expression of *TPSB2*, which is a protein produced almost exclusively by mast cells and widely used to identify them. Furthermore, genes down-expressed by *ATRX*-altered poorly differentiated sarcomas are mostly linked to adaptive immune cell activation, so adaptive immune cells are either less present or less active. This could be achieved by avoiding the release of chemoattractants and hence the recruitment or activation of other immune cells. The precise mechanism involved upon ATRX loss that changes the immune microenvironment of sarcomas needs to be deciphered.

Regarding *ATRX* expression, 27% of cases (17/63) showed no nuclear ATRX protein, which is consistent with the literature ^14,23,24,31^. In tumors presenting FS/NS, 87.5% (14/16) exhibited no ATRX protein at all. In these cases, *ATRX* mRNA level was low, likely meaning that if the truncated protein is expressed (missed by our screening with the C-terminal antibody), it should be very low. Moreover, if truncated proteins are expressed, the lost domains should be the same in all studied sarcoma types, with partial or complete loss of the helicase C-terminal domain in 90% of cases (18/20) and of both helicase domains in 70% of cases (14/20). As the majority of MS mutations occurred in one helicase domain (71.4%, 5/7) and IHC detected a nuclear localization of the protein, a decrease in ATRX enzymatic activity may be hypothesized ^32^. Collectively, these results suggest that alterations of *ATRX* preferentially target its enzymatic functions rather than its protein-protein interactions, thus explaining why mutations in *ATRX* partner genes (*i.e. DAXX*, *EZH2, SP100*) are not frequent and not an alternative to *ATRX* alteration in sarcomas. We thus hypothesize that, by modifying its chromatin remodeling action, alterations of *ATRX* trigger a specific transcriptomic program that promotes attenuated mast cell recruitment, leading to the observed immune response in models and human tumors.

Our findings show that *ATRX* alterations are quite frequent in pleomorphic sarcomas (close to 25%) and mostly lead to the loss of ATRX. In addition, we demonstrate that *ATRX* alterations are not only associated with ALT phenotype but also with differentiation and immune response regulation through non-recruitment of mast cells. Currently, most immunotherapies of sarcomas, which target the adaptive immune system and specifically T cells by helping them to recognize tumors, have a low response rate ^33^. Indeed, several recent trials have assessed the response to checkpoint inhibitors, which are used to thwart immune system escape by activating CD8+ cytotoxic T cells, with an overall response rate not higher than 25% ^34,35^. A better response rate with around 66% of response was reached by transferring autologous T cells transduced with a T cell receptor directed against a cancer antigen (CAR-T cell) expressed by the selected tumors, but only 6 patients were involved ^36^. As targeting the adaptive immune system does not work well in sarcomas, some have tried to target the innate immune system by making therapeutic vaccines which rely on the activation of dendritic cells in the presence of predetermined immunogenic antigen ^33^. One trial presented 10 out of 23 patients who lived more than 1 year whereas others died after around 7 months ^37^ and another one showed a 1-year progression-free survival of 70.6% ^38^. Targeting the innate system might therefore lead to a better outcome for sarcoma patients that could be further improved by assessing *ATRX* status before testing mast cell-enhancing therapies, as they have been successful in other solid tumors ^39^. These therapies enhanced local mast cells degranulation by using IgE antibodies, as proposed by Singer and Jensen-Jarolim ^40^. This strategy could be useful in *ATRX^WT^* tumors to enhance the anti-tumoral action of mast cells and, in *ATRX*-altered sarcomas, to enhance mast cell recruitment and activation ^41^.

## Acknowledgments

The authors would like to thank the Centre Nacional d’Anàlisi Genòmica (CNAG, Barcelona, Spain) for WG and RNA sequencing services. We acknowledge the personnel of CREFRE US006 and Animal Facility A2 for their technical assistance. We thank Mickaël Michaud for his technical and theoretical advice and Michel Charbonneau and Nathalie Grandin for their work on ALT detection in cell lines. We acknowledge Françoise Redini for providing us MG63 and K7M2 cell lines.

## Authors’ Contributions

Conceptualization, L.D, G.P, N.T, F.C; Methodology, L.D, G.P, N.T, F.C; Software N.T, N.D, L.G, T.L, E.D; Validation, L.D, G.P; Formal analysis, L.D, G.P, N.T, F.C; Investigation, L.D, G.P, S.LG, L.D, A.G-B, E.R, J.B, L.L; Writing – Original Draft, L.D, G.P, N.T, F.C; Writing – Review & Editing, L.D, G.P, N.T, F.C, L.D, T.L, E.D; Visualization: L.D, G.P, N.T; Supervision: F.C; Funding Acquisition: J.M.C, J.Y.B, F.C, Other (data and expertise): The French Sarcoma Group and the International Cancer Genome Consortium

## Methods

Further information and requests for resources should be directed to Frédéric Chibon

### Data Availability

Cohort 1 ICGC Whole-Genome sequencing and RNA sequencing data for the 67 LMS are available at https://dcc.icgc.org/projects/LMS-FR.

Cohort 2 RNA-seq expressions are available on Gene Expression Omnibus under accession GSE71121. RNA-seq raw files (FastQ) are available on sequence read archive under accessions SRP059588 or SRP059588.

Mouse K7M2 tumors expression data are available on Gene Expression Omnibus (GEO) under accession GSE157953 and will be publicly released on 09/01/2021 and available before by contacting the Lead Contact.

### Human samples

Samples used in cohort 1 were collected prospectively by the French Sarcoma Group as part of the ICGC program (International Cancer Genome Consortium). Samples used in cohort 2 were part of the cohort used in Lesluyes *et al.* ^18^. Clinico-pathological data and patient information are summarized in Table S1. All cases were systematically reviewed by expert pathologists of the French Sarcoma Group according to the World Health Organization guidelines ^42^.

From those samples, DNA extraction, whole genome sequencing and analysis, RNA extraction, sequencing and analysis as well as, annotation of variants and break point detection can be found in supplementary information.

### Validation of ATRX alterations

For cohort 1, all FS were verified at both DNA and RNA levels by Sanger sequencing. All SV were verified on DNA by Sanger sequencing and the effect on RNA was detected by RNAseq. Regarding MS and NS mutations, only those not found in both WG-seq and RNAseq were verified by Sanger sequencing. Complete deletion of *ATRX* due to total loss of chrXq or chrX was seen in 4 females all presenting RNA expression and nuclear protein, implying that the loss occurred on the inactive X. One triploid tumor developed in a male also presented a deletion of the gene but with one copy left. Therefore, they were all considered as WT regarding *ATRX* alteration. For cohort 2, whole *ATRX* cDNA was sequenced by Sanger sequencing for all cases and alterations found at RNA level were verified on DNA.

### PCR on genomic DNA

For screening of mutations on genomic DNA, PCR primers were designed using the Primer 3 program ^55^(https://bioinfo.ut.ee/primer3-0.4.0/) and are presented in Table S5. All PCR were performed on 50ng of DNA using AmpliTaqGold^®^ DNA polymerase (4311820, Applied Biosystems, Foster City, CA, USA) according to the manufacturer’s instructions with the PCR program described in the Table S5 Legend.

### RT-PCR

Total RNA was first reverse-transcribed using random hexamers and the High Capacity cDNA Reverse Transcription Kit (4368814, Applied Biosystems, Foster City, CA, USA) according to the manufacturer’s instructions. All primers used were designed using the Primer 3 program ^55^(https://bioinfo.ut.ee/primer3-0.4.0/). For *ATRX* cDNA screening, primers used are presented in Table S6. For fusion transcript detection, control PCR were first performed with different forward and reverse primers for each gene implicated in the fusion, and then PCR was performed using a forward primer for one gene and reverse primer for the other gene (Table S7). All PCR were performed as previously described for PCR on genomic DNA.

### Sanger Sequencing

PCR products were purified using an ExoSAP-IT PCR Purification Kit (US78200, GE Healthcare, Piscataway, NJ, USA) and sequencing reactions were performed with the Big Dye Terminator V1.1 Kit (4336805, Applied Biosystems, Foster City, CA, USA) according to the manufacturer’s recommendations. Samples were purified using the Big Dye XTerminator Purification kit (4376486, Applied Biosystems, Foster City, CA, USA) according to the manufacturer’s instructions and sequencing was performed on a 3730xl Genetic Analyzer for cohort 1 or 3130xl Genetic Analyzer for cohort 2 (Applied Biosystems, Foster City, CA, USA). Sequences were then analyzed using the Sequencing analysis V5.3.1 and the SeqScape V2.6 software (Life Technologies, Carlsbad, CA, USA). FinchTV software (V1.4.0) was also used (Geospiza, Seattle, WA, USA).

### Immunohistochemistry

Sixty-seven tumors in cohort 1 were analyzed on tissue microarrays. Each case was represented by three spots 4-μm-thick and 1mm in diameter. Immunohistochemistry was performed on a BenchMark Ultra instrument (Ventana, Washington D.C, USA). Antigen retrieval was performed using a CC1 protocol for 16 min at 98°C (Ventana, Washington D.C, USA), and the anti-ATRX antibody (1:1000, BSB3297, Clone BSB-108, Diagomics, Blagnac, France) was diluted in PREPKIT9 for 20 min. Antibody detection was performed using the Optiview detection kit (860-099, Ventana, Washington D.C, USA). Immunohistochemical pictures were taken using a Panoramic 250 Flash II Digital Slide Scanner and analyzed with the Panoramic Viewer (3DHISTECH Ltd., Budapest, Hungary).

Immunolabeling for ATRX was considered as positive if tumor cells had nuclear labeling, whatever its intensity (1, 2 or 3), with no evidence of cytoplasmic labeling. Neoplasms were scored as negative for ATRX if there was no labeling. One tumor presenting cytoplasmic sequestration with a strong intensity was considered as interpretable. The internal controls (inflammatory and endothelial cells) had to be positive with a nuclear labeling; otherwise the case was considered as not interpretable.

### Immunofluorescence

One hundred and twenty-seven tumors from the two cohorts were analyzed on tissue microarrays. Tissues were deparaffinized in xylene and rehydrated in a series of ethanol baths. For antigen retrieval, slides were incubated in DAKO Target Retrieval Solution, pH 9 (S236784-2, DAKO, Carpinteria, CA, USA), for 20 min in a microwave oven. The primary antibodies and dilutions (dilution in DAKO REAL antibody diluent, S202230-2, DAKO, Carpinteria, CA, USA) used to study ALT were as follows: anti-PML (1:200, PG-M3, RRID:AB_628162, sc-966, Santa-Cruz, Dallas, TX, USA) and anti-TERF2 antibody (1:200, HPA001907, RRID:AB_1080246, Sigma, St Louis, MO, USA). All primary antibodies were incubated for 1h at room temperature (RT). Secondary antibodies and dilutions used were as follows: anti-Mouse Immunoglobulins/FITC (1:400, F0479, Dako, Carpinteria, CA, USA) and anti-Rabbit IgG (H+L) Alexa Fluor® 594 conjugate (1:500, A-11072, RRID:AB_2534116, Thermo Fischer Scientific, Waltham, MA, USA). Slides were mounted with Vectashield/DAPI medium (H-1200-10, Vector Laboratories, Burlingame, CA, USA) and were then analyzed under a Nikon Eclipse 80i (Nikon, Melville, NY, USA) fluorescent microscope with appropriate filters. Pictures were captured using a Hamamatsu C4742-95 CCD camera (Hamamatsu, Hamamatsu City, Japan).

To study tryptase, tissue sections were blocked with 5% mouse serum PBS1X for 1h30 and incubated with mouse anti-tryptase antibody (1:300, ab2378, RRID:AB_303023, ABcam, Cambridge, UK) for 1h at RT. Then Alexa Fluor Plus 594 goat anti-mouse secondary antibody (1:400, A-32742, RRID:AB_2762825, Thermo Fischer Scientific, Waltham, MA, USA) was incubated for 1h at RT. Slides were mounted using the Vectashield mounting medium plus DAPI (H-1200-10, Vector Laboratories, Burlingame, CA, USA). Images were acquired on a Zeiss Cell Observer microscope (Carl Zeiss, Oberkochen, Germany). Percentage of mast cells was assessed by counting the number of mast cells in 10 same size randomly localized regions of interest (ROI) in each tumor, divided by the total number of cells in these ROI determined by the number of nuclei count with Fiji ^56^.

### Cell lines and primary culture

The cell lines MG63 (RRID:CVCL_0426; Male) and K7M2 (RRID:CVCL_V455; Female) were given by Dr. Françoise Redini. Those and HEK293T (RRID:CVCL_0063; Female) cells were cultured in DMEM (31966-021, Life Technologies, Carlsbad, CA, USA). IB106 (UPS; Female) cell is a primary culture established as previously described ^43^ and was cultured in RPMI-1640 (524000-025, Life Technologies, Carlsbad, CA, USA). Both medium were supplied with 10% fetal bovine serum (S1810-500, Dutscher, Brumath, France) and cells were kept at 37°C in a humidified chamber containing 5% CO2.

### ShRNA Knockdown of ATRX expression

shRNAs constructs targeting human or mouse *ATRX* were obtained from OriGene (Rockville, MD, USA). The 28 bp human sequence was 5’- CCTTCTAACTACCAGCAGTTGATATGAG -3’ (TL306482A, OriGene, Rockville, MD, USA) and the 29 bp mouse sequence was 5’- CATCAAGTAGATGGTGTTCAGTTTATGTG -3’ (TL502431B, OriGene, Rockville, MD, USA). A shRNA 29-mer scrambled shRNA was used as a negative control (TR30021V, OriGene, Rockville, MD, USA).

### Production of lentiviruses

Lentiviruses were produced by co-transfection of pVSVg (RRID:Addgene_138479), psPAX2 (RRID:Addgene_12260) and shRNA construct in HEK293T cells. Co-transfection was performed by adding these plasmids, chloroquine at 0.025 mM (C6628, Sigma, St Louis, MO, USA), CaCl2 at 0.125 M (C5050, Sigma, St Louis, MO, USA) and HeBS 1X (51558, Sigma, St Louis, MO, USA), HEK293T cells were then incubated at 37°C in a humidified chamber containing 5% CO2. After 6 hours, HEK293T cell medium was changed with RPMI-1640 (524000-025, Life Technologies, Carlsbad, CA, USA) containing 10% of fetal bovine serum (S1810-500, Dutscher, Brumath, France).

### Lentiviral transduction

HEK293T cell culture medium was filtered with a 0.45 μm PES filter and was mixed at 1:1 ratio with K7M2, MG63 or IB106 culture medium previously seeded. Polybrene (8 μg/ml, H9268, Sigma, St Louis, MO, USA) was also added with the virus. Infected cells were selected with puromycin and cells were sorted by FACS (BDFACSAria, BD Biosciences, San Jose, CA, USA) thanks to their GFP expression when vector with shRNA *ATRX* was integrated.

### Western Blot analysis

Protein extracts were separated from each cell line with RIPA protein lysis buffer (R0278, Sigma, St Louis, MO, USA) containing 1X protease cocktail (P8340, Sigma, St Louis, MO, USA). Protein extracts were separated by electrophoresis on acrylamide gel (456-8085, Bio-Rad, Hercules, CA, USA) and transferred onto PVDF membrane. Then they were probed with antibodies against ATRX (1:1000, HPA001906, RRID:AB_1078249, Sigma, St Louis, MO, USA) or actin (1:5000, A5316, RRID:AB_476743, Sigma, St Louis, MO, USA). Proteins of interest were detected with HRP-conjugated horse anti-mouse IgG antibody (1:5000, 7076S, RRID:AB_330924, Cell Signaling Technology, Danvers, MA, USA) or HRP-conjugated goat anti-rabbit IgG antibody (1:5000, 7074S, RRID:AB_2099233, Cell Signaling Technology, Danvers, MA, USA) and visualized with the ECL prime Western blotting detection regent (RPN2236, GE Healthcare, Piscataway, NJ, USA), according to the manufacturer’s recommendations and using the PXi system.

### ALT specific c-circle detection

The C-circle assay, which partially detects single-stranded telomeric (CCCTAA)n DNA circles (C-circles) amplified by the Phi29 polymerase in the absence of dCTP, was performed as previously described ^61^.

### Cell proliferation assay

Cells of each cell line with ATRX^KD^ or ATRX^CT^ were seeded onto a 96-well plate (3.10^3^ cells/well). After 4 days, cell proliferation was evaluated by adding 20 μL of MTT (M2128, 5mg/mL, Sigma, St Louis, MO) to cell medium. Two hours later, medium was replaced by 100 μL of DMSO (5879, Sigma, St Louis, MO, USA) and the optic density (OD) of each well was read with a spectrophotometer at 570 and 630 nm. Live cell number was correlated to Δ*OD* = *OD*_570*nm*_ − *OD*_630*nm*_. Experiments were performed independently in triplicate three times.

### Soft agar assay

Cells of each cell line with ATRX^KD^ or ATRX^CT^ were seeded (5000 cells/well) in 0.35 % agarose cell medium (16500-500, Invitrogen, Carlsbad, CA, USA) onto a 6-well plate containing a 0.5 % agar base. 0.5 mL of cell culture medium was added and changed every 3-4 days. After incubating for 3 to 4 weeks, colonies were visualized with 0.005 % crystal violet staining (HT90132, Sigma, St Louis, MO, USA). Experiments were performed independently in triplicate four times.

### In vivo experiment

All experiments were performed in conformity with the rules of the French Institutional Animal Care and Use committee (approval number DAP-APAFiS-2018041617309605) and all efforts were made to minimize animal suffering. Mice were maintained under specific pathogen-free conditions in the animal facility of University of Bordeaux (France) or at the CREFRE (Centre Régional d’Exploration Fonctionnelle et Ressources Expérimentales, Toulouse, France).

For the experiment with IB106 cells, 6-8-week-old female NSG (NOD.Cg-Prkdc^scid^ Il2rg^tm1Wjl^/SzJ; RRID:BCBC_4142) mice were used. Ten mice were injected with 800,000 IB106 ATRX^KD^ cells and ten with 800,000 IB106 ATRX^CT^ cells as controls. Mice were randomly assigned to one cage of five animals then each cage was randomly assigned to a group of cells. One mice in the control group was exclude due to an important and fast loss of weight so the final number of unit is N=10 in the ATRX^KD^ group and N=9 in the ATRX^CT^ one. Tumor sizes were measured without knowing the affiliation group of each mice twice a week using a caliper and their volume was calculated using the formula: (L^2^×l)⁄2. At the end of the experiment, mice were sacrificed by cervical dislocation. Tumors were then weighed and divided in two parts for formalin fixation and nitrogen freezing. Each tumor was stained with HE and analyzed by a pathologist specialized in sarcomas. Growth rate was calculated with the segmental linear regression of GraphPad Prism (GraphPad Software, San Diego, CA, USA) and statistical analyses were done using an unpaired T-test.

For the experiments with K7M2 cells, 6-8 weeks-old female NSG (NOD.Cg-Prkdc^scid^ Il2rg^tm1Wjl^/SzJ; RRID:BCBC_4142) or Balb/c (Balb/cJ; RRID:IMSR_JAX:000651) mice were used. For groups were made, each group was composed by 15 mice, one group of NSG mice was injected with 500,000 K7M2 ATRX^KD^ cells, the other with 500,000 K7M2 ATRX^CT^ cells as controls. The Balb/c mice were injected in the same conditions. Mice were randomly assigned to one cage of five animals then each cage was randomly assigned to a group of cells. One animal from the Balb/c ATRX^CT^ group was exclude due to a teeth malformation and an incapacity to eat so the final number of unit is N=15 in each group except this one which was N=14. Tumor sizes were measured without knowing the affiliation group of each mice twice a week using a caliper and their volume was calculated using the formula: (L^2^×l)⁄2. At the end of the experiment, mice were sacrificed by cervical dislocation. Tumors were then weighed and divided in two parts for formalin fixation and nitrogen freezing. Each tumor was stained with HE and analyzed by a pathologist specialized in sarcomas. Growth rate was calculated with the segmental linear regression of GraphPad Prism (GraphPad Software, San Diego, CA, USA) and statistical analyses were done using an unpaired T-test. Survival curves were analyzed with GraphPad Prism using the Kaplan-Meier method.

### Mice tumor RNA sequencing and analysis

Total RNA was extracted, prepared and sequenced as described above to obtain more than 20 million paired-end reads with a length of 75 bp each. Bioinformatic analysis was done as previously described ^18^.

RNA reads were aligned to the mm10 genome assembly with STAR v2.6.0c ^57^. Low-quality (score < 20) and duplicated PCR paired-end reads were removed with SAMtools v1.8 ^46^ and PicardTools v2.18.2 ^47^(http://broadinstitute.github.io/picard/), respectively. Then, gene expression was quantified with Cufflinks v2.2.1 ^58^, using RefSeq ^59^ genes (without miRNA and rRNA) from mm10 UCSC Table Browser ^60^ fixed on 2019/01.

Differential gene expression was performed by R package DESeq, between ATRX^KD^ and ATRX^CT^ tumors extracted from Balb/c mice. Relationships between proteins overexpressed in ATRX^KD^ and ATRX^CT^ tumors were assessed by the STRING Database ^19^.

### Quantification and statistical analysis

Kaplan-Meier analyses were performed for metastasis-free survival and overall survival. To subdivide *ATRX* expression in two groups, expression was plotted for *ATRX* WT and altered cases, separately. The intersection between these two density curves was 4.45 (log2 FPKM) and 2.77 for cohort 1 and 2, respectively.

Differential gene expression (DGE) analyses were performed by R package DESeq. Gene Ontology (GO) analysis was performed on these differentially expressed genes (P < 0.05 and fold-change > 2 or < −2), by R package GOseq. In parallel, significant genes with P < 0.01 were used to make a heatmap (R package ComplexHeatMap).

Every other statistical analyses details can be found in each figure legend.

