## Supplemental Methods/Figures/Tables for "ATRX alteration contributes to tumor growth and immune escape in pleomorphic sarcomas"

### Supplementary figures and legends:

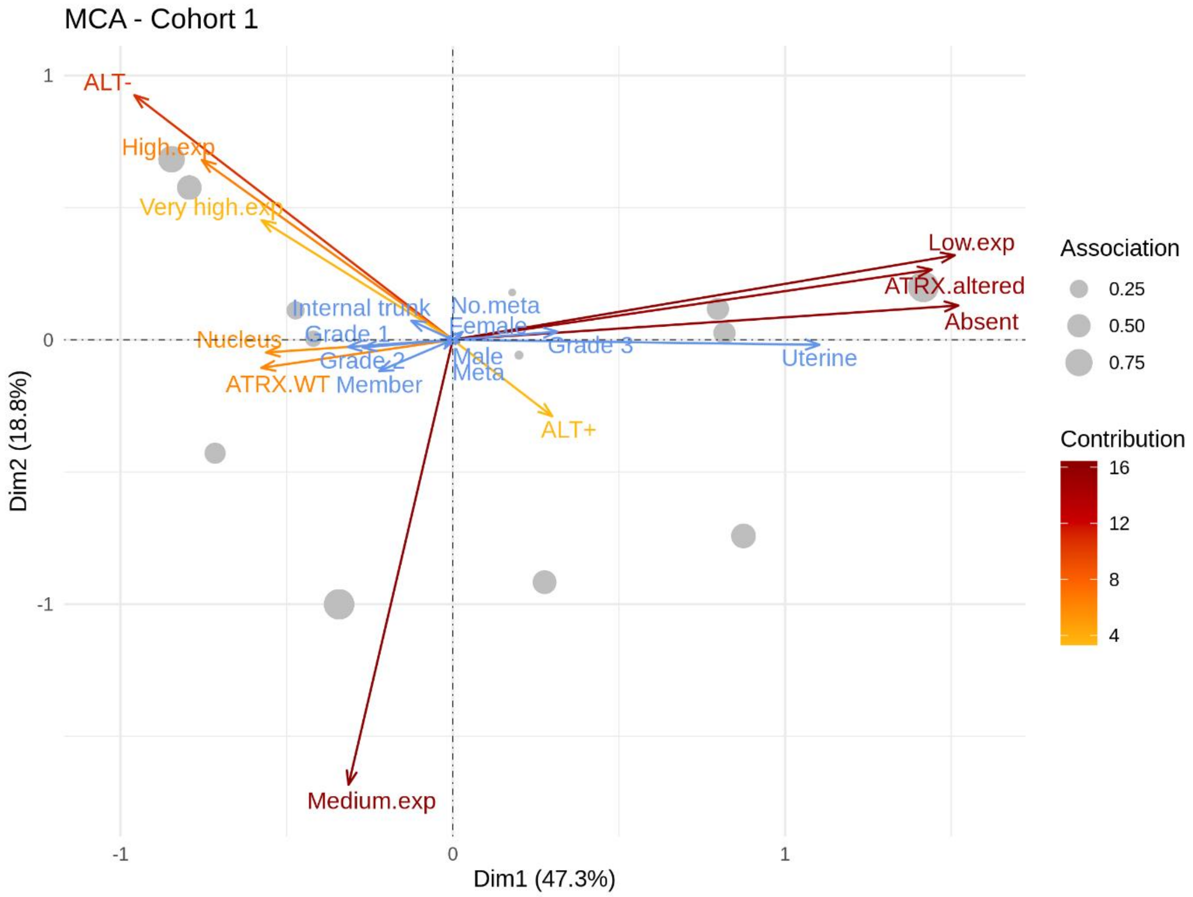


**Figure S1**: Multiple correspondence analysis related to Figure 2

Multiple correspondence analysis (MCA) was performed on *ATRX* status, mRNA expression, ATRX location and ALT phenotype. Additional variables and sample groups are represented in blue arrows and gray points, respectively.

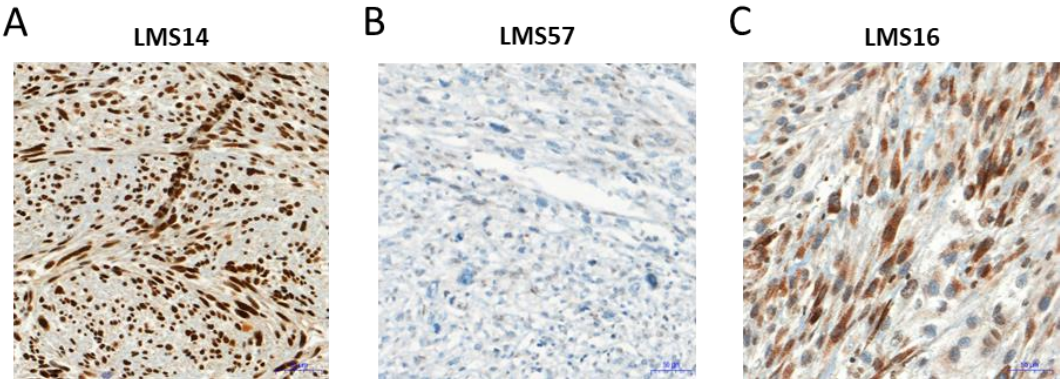


**Figure S2**: Example of ATRX staining by immunohistochemistry related to Figure 2

Leiomyosarcomas with **A** strong nuclear staining, **B** no tumorous staining or **C** cytoplasmic staining. In all samples, non-tumorous cells serve as internal control and have nuclear staining.

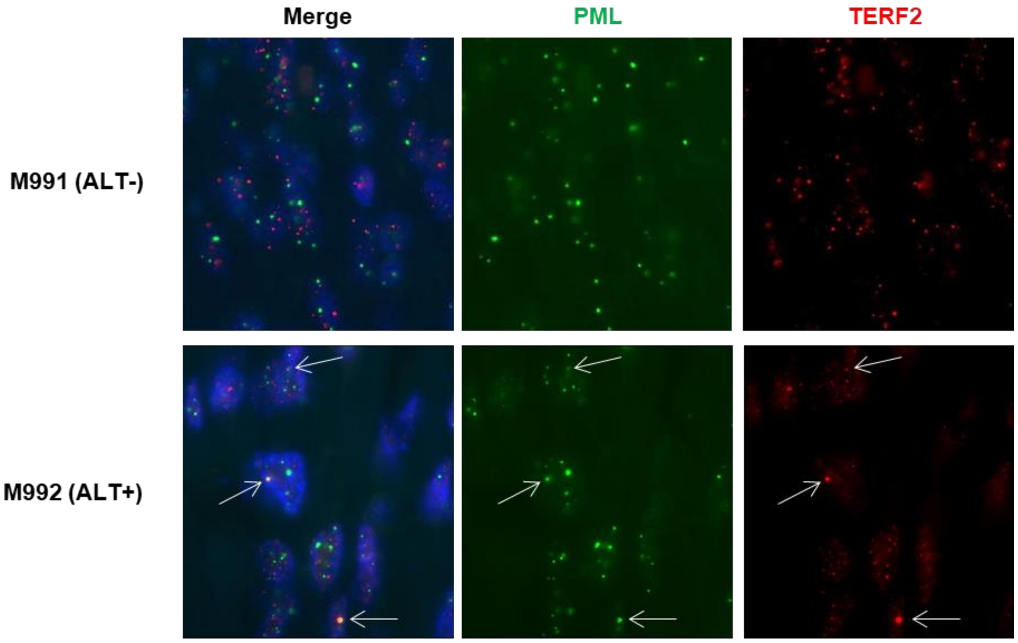


**Figure S3**: ALT status determined by PML/TERF2 immunofluorescence analysis related to Figure 2

Immunofluorescence showing absence of co-localization of PML (green) and TERF2 (red) signals in a tumor sample (ALT-) and of PML and TERF2 protein co-localization (ALT+), indicating that the ALT mechanism should be active in this tumor (ALT+). Protein co-localizations are highlighted by arrows. Magnification: X1000.

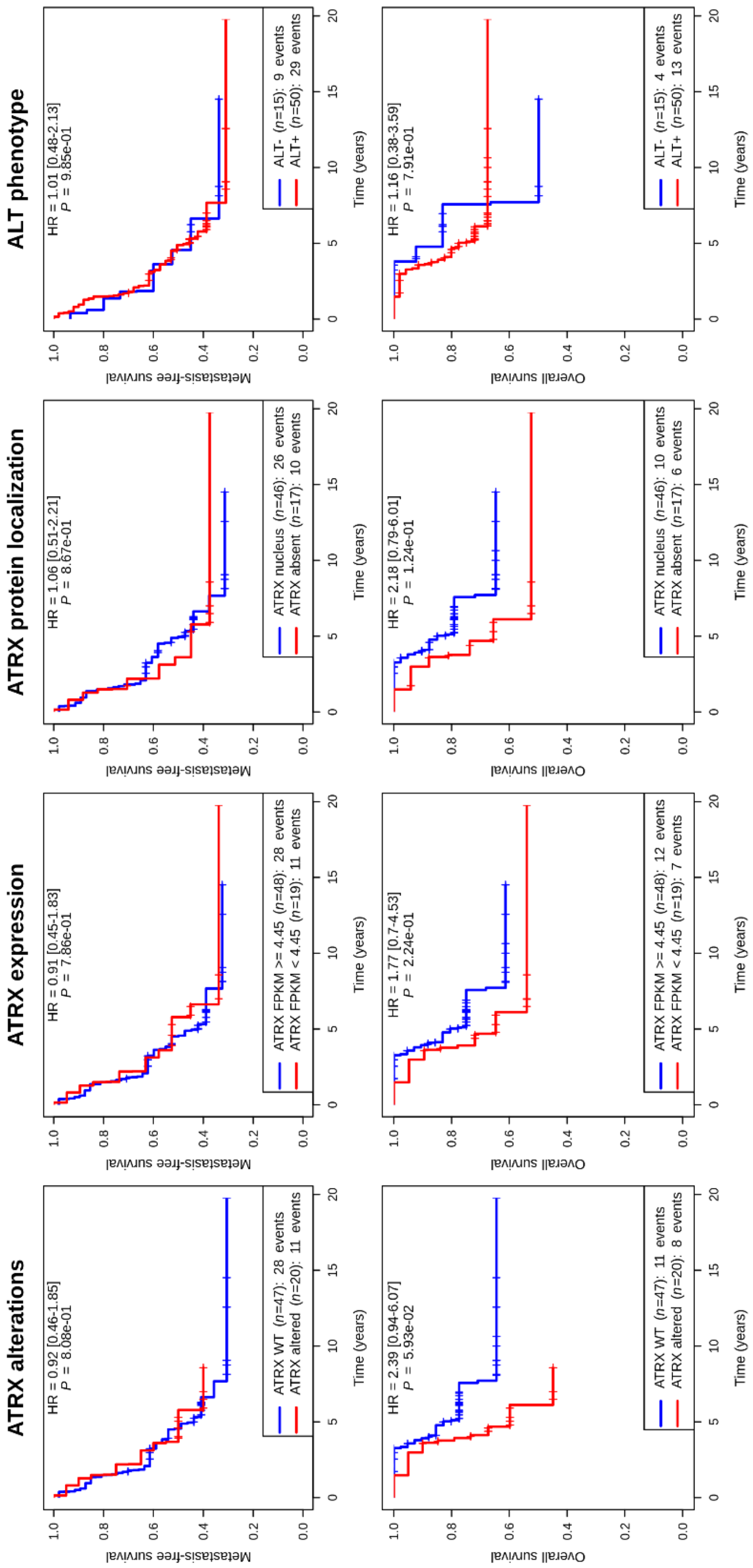


**Figure S4**: Kaplan-Meier analysis in leiomyosarcomas (Cohort 1), according to *ATRX* status (wild-type vs altered), *ATRX* expression (high *vs* low), ATRX localization (nucleus *vs* absent) and ALT mechanism phenotype (ALT- *vs* ALT+). Up: metastasis-free survival. Down: overall survival. To subdivide *ATRX* expression into two groups, it is plotted for *ATRX* wild-type and altered cases, separately. Intersection between these two density curves is 4.45 (log2(FPKM+1)) related to Figure 2.

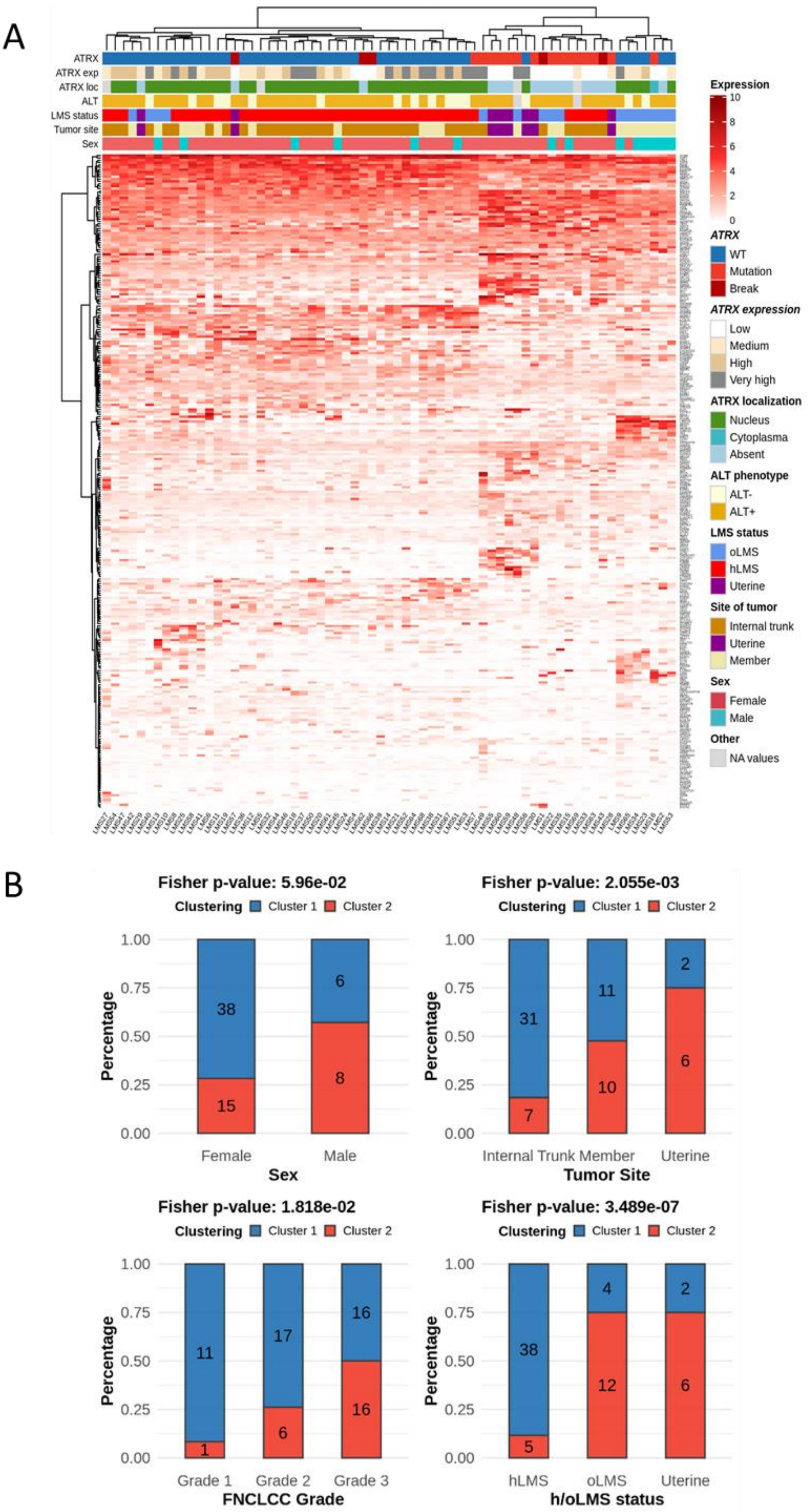


**Figure S5**: Clustering with differentially expressed genes according to ATRX status related to Figure 3

**A** mRNA expression (log2(FPKM+1)) of 279 differentially expressed genes in *ATRX* altered tumors (see also Figure 3A; $P \leq 0.01$ and fold-change $\leq -2$ or $\geq 2$). **B** Association between cluster 1 or 2 (left and right cluster, respectively) to sex, tumor site, grade and LMS status, oLMS and hLMS means “other” and “homogeneous” LMS, respectively (Darbo *et al*, 2020).

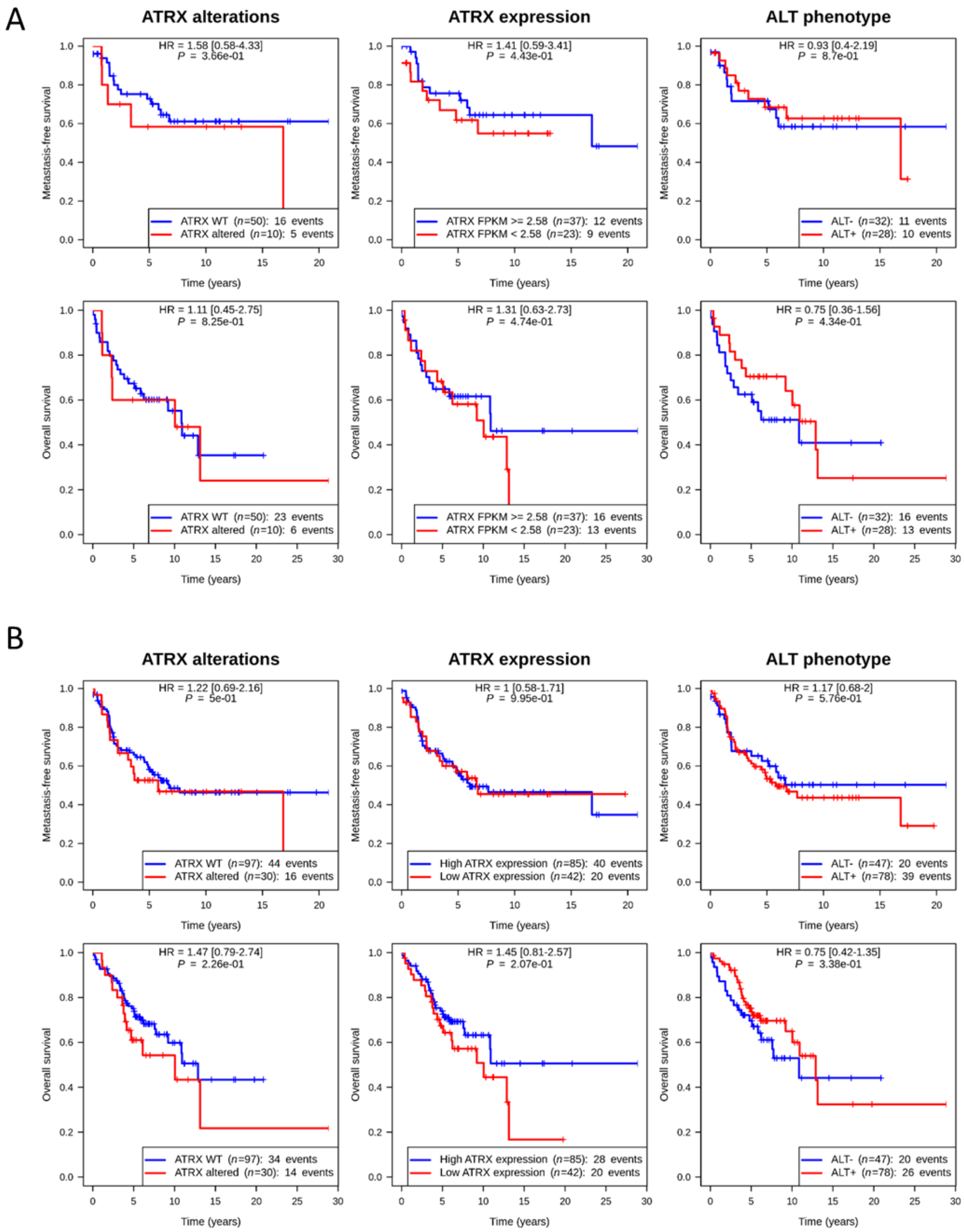


**Figure S6**: Kaplan-Meier analyses related to Figure 4.

**A** Kaplan-Meier analysis in poorly differentiated sarcomas (Cohort 2) according to *ATRX* status (wild-type *vs* altered), *ATRX* expression (high *vs* low) and ALT phenotype (ALT- *vs* ALT+). Up: metastasis-free survival. Down: overall survival. To subdivide *ATRX* expression into two groups, it is plotted for *ATRX* wild-type and altered cases separately. Intersection between these two density curves is 2.58 (log2(FPKM+1)). **B** Kaplan-Meier analysis in leiomyosarcomas and poorly differentiated sarcomas (Cohorts 1 and 2), depending on *ATRX* status (wild-type *vs* altered), *ATRX* expression (high *vs* low) and ALT mechanism phenotype (ALT- *vs* ALT+). Up: metastasis-free survival. Down: overall survival. To subdivide *ATRX* expression into two groups, the two cohorts are taken separately to identify the cut-off (see also Supp. Figures 4A and 6A).

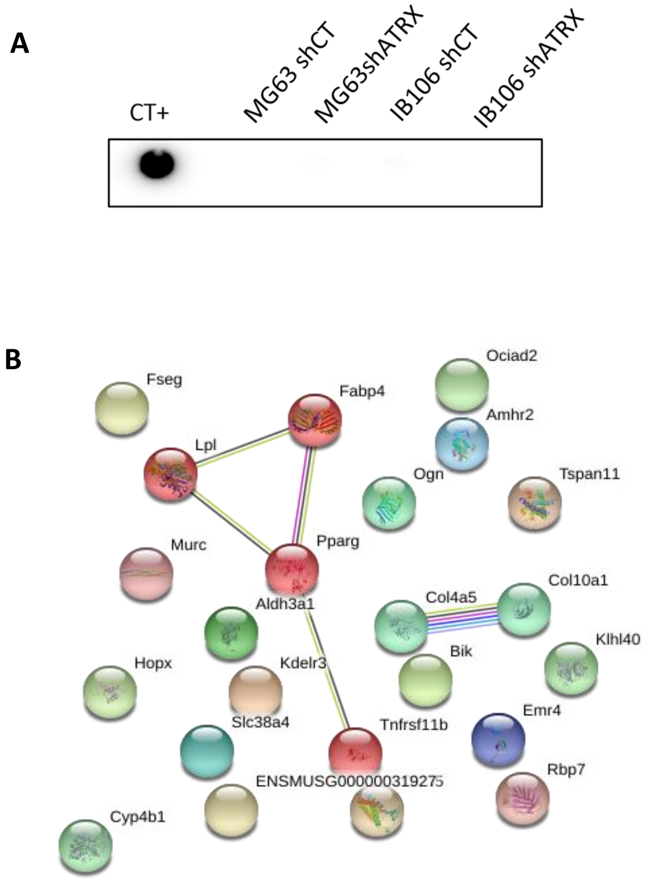


**Figure S7**: shRNA against ATRX effect on sarcoma cell lines related to Figure 6 and 7.

**A** C-circle analysis of ALT status upon shRNA against *ATRX* or in control condition. Positive control is U2OS DNA. **B** Links between proteins encoded by genes overexpressed in K7M2 $ATRX^{KD}$ tumors displayed by STRING Database showing two clusters via MCL cluster.

### Supplementary tables and legends:

| | Cohort1 (n=67) | Cohort2 (n=60) |
|---|---|---|
| **FOLLOW-UP and OUTCOMES** | | |
| Follow-up (years) | | |
| Median | 5.22 | 5.94 |
| Range | 1.48-19.77 | 0.01-28.88 |
| **PATIENT CHARACTERISTICS** | | |
| Age at diagnosis (years) | | |
| Mean | 62.94 | 62.95 |
| Median | 64 | 64.5 |
| Range | 22-80 | 20-87 |
| Gender | | |
| Female | 53 (78.46%) | 22 (36.67%) |
| Male | 14 (21.54%) | 38 (63.33%) |
| **TUMOR CHARACTERISTICS** | | |
| Tumor site | | |
| Internal trunk | 38 | 9 |
| Uterine | 8 | 0 |
| Member and Trunk wall | 21 | 51 |
| Tumor depth | | |
| Deep | 56 | 42 |
| Superficial | 7 | 5 |
| Superficial and deep | 4 | 13 |
| Tumor size (cm) | | |
| Median | 8 | 8 |
| Range | 1.5-23 | 1-30 |
| Histotype | | |
| LMS | 67 | 0 |
| UPS | 0 | 30 |
| MFS | 0 | 17 |
| DDLPS | 0 | 13 |
| FNCLCC Grade | | |
| I | 12 | 4 |
| II | 23 | 18 |
| III | 32 | 40 |
| Resection status, margins | | |
| R0 | 42 | 24 |
| R1 | 18 | 28 |
| R2 | 1 | 2 |
| Unknown | 6 | 6 |

**Table S1:** Clinical characteristics in both cohorts related to Figure 1, 2 and 4.

LMS: leiomyosarcoma, UPS: undifferentiated pleomorphic sarcoma, MFS: myxofibrosarcoma, DDLPS: dedifferentiated liposarcoma.

| Cohort | Sample | Histotype | Location | Sex | Allele 1 | Allele 2 | DNA (WGseq) Reads alt/total | DNA (WGseq) Mut. allele freq. | RNA (RNAseq) Reads alt/total | RNA (RNAseq) Mut. exp. freq. | RNA (RNAseq) *ATRX*: log2 (FPKM+1) | Functional *ATRX* status | gDNA (hg19) | cDNA (NM_000489.6) | Protein |
|---|---|---|---|---|---|---|---|---|---|---|---|---|---|---|---|
| 1 | LMS16T | LMS | Member | M | Frameshift | Y | 8/12 | 0,67 | 25/34 | 0,74 | 3,61 | 0 | chrX: g.76874277_76874281del | c.5441_5445del | p.(C1814SfsX45) |
| 1 | LMS22T | LMS | Member | M | Frameshift | Y | 11/17 | 0,65 | 18/36 | 0,50 | 4,10 | 0 | chrX: g.76875903_76875923delinsAAAT | c.5212_5232delinsTTTA | p.(I1738FfsX11) |
| 1 | LMS35T | LMS | Member | F | Frameshift | | 21/59 | 0,36 | 19/46 | 0,41 | 4,06 | 1/2 | chrX: g.76939939dupG | c.807dupC(;)485_487del | p.(E271RfsX11)(;)(E162del) |
| 1 | LMS33T | LMS | Internal Trunk | F | Frameshift | | 32/61 | 0,52 | 109/131 | 0,83 | 4,07 | 0 | chrX: g.76938713_76938734del | c.2014_2035del | p.(P672KfsX17) |
| 1 | LMS63T | LMS | Internal Trunk | F | Frameshift | | 28/52 | 0,54 | 39/56 | 0,70 | 3,99 | 0 | chrX: g.76938581_76938618dup | c.2130_2167dup | p.(E723VfsX26) |
| 1 | LMS59T | LMS | Uterus | F | Frameshift | | 12/52 | 0,23 | 34/37 | 0,92 | 3,09 | 0 | chrX: g.76938870delT | c.1878delA | p.(K626NfsX23) |
| 1 | LMS60T | LMS | Uterus | F | Frameshift | | 30/54 | 0,56 | 4/5 | 0,80 | 3,84 | 0 | chrX: g.76849230delG | c.6046delC | p.(H2016NfsX34) |
| 1 | LMS15T | LMS | Internal Trunk | M | Nonsense | Y | 22/27 | 0,81 | 18/25 | 0,72 | 3,86 | 0 | chrX: g.76778786C>A | c.6793G>T | p.(E2265X) |
| 1 | LMS28T | LMS | Uterus | F | Nonsense | | 17/47 | 0,36 | 44/48 | 0,92 | 4,42 | 0 | chrX: g.76889056C>A | c.4954G>T | p.(E1652X) |
| 1 | LMS30T | LMS | Uterus | F | Nonsense | | 20/44 | 0,45 | 34/38 | 0,89 | 4,14 | 0 | chrX: g.76938583G>C | c.2165C>G | p.(S722X) |
| 1 | LMS55T | LMS | Uterus | F | Nonsense | | 18/44 | 0,41 | 18/18 | 1,00 | 4,07 | 0 | chrX: g.76907651G>A | c.4510C>T | p.(R1504X) |
| 1 | LMS48T | LMS | Member | F | Missense | | 32/67 | 0,48 | 0/66 | 0,00 | 5,23 | 1 | chrX: g.76849167C>T | c.6109G>A | p.(V2037I) |
| 1 | LMS49T | LMS | Internal Trunk | F | Missense | Isodisomy | 41/45 | 0,91 | 118/125 | 0,94 | 5,51 | 0 | chrX: g.76875980C>T | c.5155G>A | p.(D1719N) |
| 1 | LMS69T | LMS | Internal Trunk | F | Missense | Loss | 23/28 | 0,82 | 59/61 | 0,97 | 4,49 | 0 | chrX: g.76890104C>T | c.4790G>A | p.(G1597D) |
| 1 | LMS7T | LMS | Internal Trunk | F | Missense | | 18/51 | 0,35 | 95/111 | 0,86 | 5,19 | 0 | chrX: g.76890113T>C | c.4781A>G | p.(H1594R) |
| 1 | LMS57T | LMS | Uterus | F | Del. & FS | | NA | NA | 39/49 | 0,80 | 4,36 | 0 | chrX: g.76882435_76982612del | c.21_4956del | p.(S7RfsX3) |
| 1 | LMS66T | LMS | Internal Trunk | F | Del. & FS | | NA | NA | 45/48 | 0,94 | 3,85 | 0 | chrX:g.76756028_76920645delinsATTCCA | c.3810_7479delins3809+1_3809+187 | p.(I1271YfsX62) |
| 1 | LMS1T | LMS | Internal Trunk | F | Del. Inv. Ins. | | NA | NA | NA | NA | 2,55 | 0 | chrX: g.76961353_77042805delins77042774_77042825inv | NA | NA |
| 1 | LMS43T | LMS | Internal Trunk | F | Del. | | NA | NA | NA | NA | 1,91 | 0 | chrX: g.77000959_77060888del | NA | NA |
| 1 | LMS62T | LMS | Internal Trunk | F | Trans. Complex | | NA | NA | NA | NA | 2,22 | 0 | Complex | NA | NA |

| Cohort | Sample | Histotype | Location | Sex | Allele 1 | Allele 2 | Sanger DNA allelic status | Reads alt/total | Mut. exp. freq. | *ATRX*: log2 (FPKM+1) | Functional *ATRX* status | gDNA (hg19) | cDNA (NM_000489.6) | Protein |
|---|---|---|---|---|---|---|---|---|---|---|---|---|---|---|
| 2 | S873 | MFS | Member | M | Frameshift | Y | B | 0/4* | Sanger: B | 2,39 | 0 | g.76874280_76874290del | c.5432_5442del | p.(L1811CfsX46) |
| 2 | N003 | DDLPS | Internal Trunk | M | Nonsense | Y | A/B (heterogenous) | 0/3* | Sanger: A/B | 0,86 | 0 | g.76938583G>C | c.2165C>G | p.(Ser722X) |
| 2 | S859 | UPS | Member | M | Nonsense | Y | B | 8/8 | 1,00 | 2,64 | 0 | g.76875866C>A | c.5269G>T | p.(E1757X) |
| 2 | S825 | UPS | Trunk wall | M | Missense | Y | B | 21/21 | 1,00 | 3,24 | 0 | g.76938098C>T | c.2650G>A | p.(E884K) |
| 2 | S928 | UPS | Member | F | Missense | | A/B | 8/8 | 1,00 | 2,67 | 0 | g.76875938T>C | c.5197A>G | p.(K1733E) |
| 2 | S832 | MFS | Member | F | Missense | | A/B | 6/6 | 1,00 | 2,90 | 0 | g.76938571T>A | c.2177A>T | p.(D726V) |
| 2 | S823 | MFS | Member | M | Fusion transcript | Y | NA | NA | NA | 0,98 | 0 | NA | NA | p.(S7RfsX62) |
| 2 | S831 | UPS | Trunk wall | M | Fusion transcript | Y | NA | NA | NA | 2,47 | 0 | NA | NA | p.(D2234NfsX32;D2234RfsX4) |
| 2 | S918 | UPS | Member | F | Fusion transcript | | NA | NA | NA | 2,16 | 0 | NA | NA | p.(V1520DfsX54;I1521VfsX33) |
| 2 | M979 | MFS | Member | F | Fusion transcript | | NA | NA | NA | 2,04 | 0 | NA | NA | p.(R2109SfsX12) |

**Table S2:** *ATRX* mutations and structural variants in both cohorts related to Figure 2 and 4.

For each of the 30 validated alterations of *ATRX* (NM_000489.6), genomic and expression information and annotations are reported in the table. Regarding cohort 2, only gDNA Sanger sequencing data for point mutations are available. *: not detected by RNAseq but detected by Sanger sequencing on cDNA. Mutation expression frequency for these two mutations could not be determined using RNAseq, probably because of tumor heterogeneity, so allele expression in RNA was defined according to Sanger sequencing data. Alt: alternative, Freq.: frequency, Exp.: expression, Mut.: mutation, Loc.: location, M: Male, F: Female, LMS: leiomyosarcoma, DDLPS: dedifferentiated liposarcoma, MFS: myxofibrosarcoma, UPS: undifferentiated pleomorphic sarcoma, Y: chromosome Y, A: reference allele, B: alternative allele, Del: deletion, Ins: insertion, Inv: inversion, FS: Frameshift, SV: structural variant, Trans.: translocation, FPKM: Fragments Per Kilobase Million, NA: not available.

| Sample | Histotype | Sex | Gene_chromosome1 | Break location | Break position (hg19) | Gene_chromosome2 | Gene name 2 | Break location 2 | Break position 2 (hg19) | Mechanism | Consequence on *ATRX* gene | Genomic nomenclature (hg19) chrX | cDNA nomenclature (NM_000489.6) | Protein nomenclature |
|---|---|---|---|---|---|---|---|---|---|---|---|---|---|---|
| | | | ***ATRX*** | | | **Fusion partner** | | | | **Annotations** | | | | |
| LMS1T | LMS | F | X | Intron 2 | 76961352 | X | **Intergenic** | X | 77042825 | Deletion/inversion-insertion | Loss of 3' part to exon 2 | g.76961353_77042805delins77042774_77042825inv | - | - |
| LMS43T | LMS | F | X | Intron 1 | 77000958 | X | **Intergenic** | X | 77060889 | Deletion | Loss of 3' part to exon 1 | g.77000959_77060888del | - | - |
| LMS57T | LMS | F | X | Intron 19 | 76882434 | X | ***ATRX*** | Intron 1 | 76982613 | Deletion | Fusion of intron 1 to intron 19 | g.76882435_76982612del | c.21_4956del | p.(S7RfsX3) |
| LMS62T | LMS | F | X | Intron 1 | 77035789 | X | ***DMD*** | Intron 60 | 31367514 | Complex | Loss of 3' part to exon 1 | Complex | - | - |
| LMS66T | LMS | F | X | Intron 10 | 76920646 | X | **Intergenic** | X | 76756027 | Deletion | Loss of exon 11 to the end | g.76756028_76920645delinsATTCCA | c.3810_7479delins3809+1_3809+187 | p.(I1271YfsX62) |

**Table S3:** *ATRX* breakpoints in cohort 1 related to Figure 1.

Chromosomal and genomic coordinates of gene breakpoint regarding *ATRX* and its fusion partners are indicated together with the potential mechanism involved and consequence for the *ATRX* gene. Nomenclatures of the alteration and its effect on RNA are indicated when an alternative RNA product was detected by RNAseq. Predicted effect on protein is then indicated. M: Male, F: Female, LMS: leiomyosarcoma.

| | | | *ATRX* | | | Fusion partner | | | | Annotations | | |
|---|---|---|---|---|---|---|---|---|---|---|---|---|
| **Sample** | **Histotype** | **Sex** | **Gene_chromosome1** | **Break location** | **Break position (hg19)** | **Gene_chromosome2** | **Gene name 2** | **Break location 2** | **Break position 2 (hg19)** | **Mechanism** | **ORF conservation** | **ATRX Protein** |
| S823 | MFS | M | X | exon 1 | 77041470 | 15 | ***DDX11L9*** | exon 1 | 102519162 | Translocation | No | p.(S7RfsX62) |
| S831 | UPS | M | X | exon 30 | 76812922 | X | ***RNU6-974P*** | downstream | 81166026 | Eversion | No | p.(D2234NfsX32;D2234RfsX4) |
| S918 | UPS | F | X | exon 15 | 76891547 | X | ***RP1-279N11.1*** | upstream | 75878708 | Deletion | No | p.(V1520DfsX54;I1521VfsX33) |
| M979 | MFS | F | X | exon 28 | 76829715 | X | ***FTX*** | intron | 73401759 | Inversion | No | p.(R2109SfsX12) |

**Table S4:** *ATRX* fusion transcripts in cohort 2 related to Figure 4.

Chromosomal and genomic coordinates of gene breakpoint regarding *ATRX* and its fusion partners are indicated together with the potential mechanism involved and predicted effect on the protein. MFS: myxofibrosarcoma, UPS: undifferentiated pleomorphic sarcoma, M: Male, F: Female.

| Primer name | Primer sequence 5'-->3' | Location | Product size (bp) | PCR program | Ref gene | Tumor |
|---|---|---|---|---|---|---|
| ATRXgDNAex7F | ACTTGTGTCCAATATGCCATTTT | Exon 7 | 366 | TD 60°C | NM_000489.6 | LMS35T |
| ATRXgDNAex7R | AGAAGTCTTCCAAGGGCAGA | | | | | |
| ATRXgDNAex9F | GCGTAATTCTTCTGACAGTGCT | Exon 9 | 346 | TD 60°C | | N003/ LMS35T |
| ATRXgDNAex9R | AGAAGACTCAGACTGGGTTTGT | | | | | |
| ATRXgDNAex17F | TGCTGTTTCTTAGAAGTTTTGGTT | Exon 17 | 485 | TD 60°C | | M987 |
| ATRXgDNAex17R | CATTAGGACCTCTGCTCAAACA | | | | | |
| ATRXgDNAex20F | CAACGATGTCATTTTATCTTCCTG | Exon 20 | 452 | TD 60°C | | S928/S859/LMS22T |
| ATRXgDNAex20R | ACCACTCATTTATAAAGCATCTCA | | | | | |
| ATRXgDNAex21F | TGAGCATTTCATTGGGGAAT | Exon 21 | 326 | TD 60°C | | S873/LMS16T |
| ATRXgDNAex21R | GCTCAGAAAATATGTTGGGATTG | | | | | |
| ATRXgDNAex26F | CTCCCAAGTCCCCATCAGTT | Exon 26 | 377 | TD 60°C | | LMS48T/LMS60T |
| ATRXgDNAex26R | AGGAAGGAAGGAAAAGCAACA | | | | | |
| ATRXBPLMS1F1 | AACTGGCAATCAAGTCTGTGC | *ATRX* intron 2 | 268 | TD 60°C | | LMS1T |
| ATRXBPLMS1R1 | TCCAAAATTCATATGGACCAGGC | | | | | |
| ATRXBPLMS1F2 | GAGGTCGAGGTTGGAGGATG | Upstream *ATRX* gene | 153 | TD 60°C | | |
| ATRXBPLMS1R2 | CACCTCAGCCTCTCAAGCAG | | | | | |
| ATRXBPLMS43F1 | CGTCCTAGCCTCTGGTAACC | *ATRX* intron 1 | 206 | TD 60°C | | LMS43T |
| ATRXBPLMS43R1 | TGGAGTCCTATTCAGCCCTT | | | | | |
| ATRXBPLMS43F2 | AATGTCTTTCTGTGCCTGGC | Intergenic chrX | 182 | TD 60°C | | |
| ATRXBPLMS43R2 | GGCAGAAGGATCGCTTGAAC | | | | | |
| ATRXBPLMS57F1 | TTCACCGTGTTAGCCAGGAT | *ATRX* intron 19 | 390 | TD 60°C | | LMS57T |
| ATRXBPLMS57R1 | ACCACCTAAATGTTGCAATACCA | | | | | |
| ATRXBPLMS57F2 | TCCACATCCTCTCCAGCATC | *ATRX* intron 1 | 476 | TD 60°C | | |
| ATRXBPLMS57R2 | GCCTATCTGCAATTGGGTCA | | | | | |
| ATRXBPLMS62F1 | AGCCTTGCCTGTACTTCTTTG | *DMD* intron 60 | 151 | TD 60°C | | LMS62T |
| ATRXBPLMS62R1 | CATGCCTGTAGTCCCAGCT | | | | | |
| ATRXBPLMS62F2 | GGCGACAGGGTGAGAGTC | *ATRX* intron 1 | 206 | TD 60°C | | |
| ATRXBPLMS62R2 | AGCAACAAAACACCTGTAACCT | | | | | |
| ATRXBPLMS66F1 | GGGAATTGAACAATGAGAACACA | Intergenic chrX | 118 | TD 60°C | | LMS66T |
| ATRXBPLMS66R1 | GTTTGCCGCACCTACTGAC | | | | | |
| ATRXBPLMS66F2 | TGCTGTTGAATAAAACCTCTCGT | *ATRX* intron 10 | 150 | TD 60°C | | |
| ATRXBPLMS66R2 | TTTGATGCTGCATAACCTTCCA | | | | | |

**Table S5:** Genomic DNA primers used for point mutations and breakpoint validation by Sanger sequencing related to methods. Forward and reverse primers used for mutation and breakpoint (BP) validations on genomic DNA are presented. All mutations were detected by two independent techniques and only those not detected by both WGseq and RNAseq were validated by Sanger sequencing. RNA primers were used for DNA validation of some *ATRX* mutations in exon 9, see Supplementary Table 6. For each BP validation in cohort 1, the two primers used to detect the BP are in red: other primers were used to detect a potential reciprocal translocation. PCR was a touch-down 60°C program (TD 60°C) (2 cycles at a temperature of 60°C, followed by 2 cycles at 59°C, 2 cycles at 58°C, 3 cycles at 57°C, 3 cycles at 56°C, 4 cycles at 55°C, 4 cycles at 54°C, 5 cycles at 53°C and finally 10 cycles at 52°C). RefSeq annotation used to locate primers for the gene is indicated. Tumors in which each primer couple was used are indicated. Ex: exon, bp: base pair, F: forward, R: reverse.

| Primer name | Primer sequence 5'-->3' | Product size (bp) | DNA validation | RNA validation |
|---|---|---|---|---|
| ATRXcDNAF1 | ACCGCTGAGCCCATGAGT | 143 | | |
| ATRXcDNAR1 | CCACTGATTTTATCTGTGTTTTGA | | | |
| ATRXcDNAF2 | TTCCTTGCACACTCATCAGAA | 399 | | |
| ATRXcDNAR2 | TTTCATTTTACTTCTGCTTCTAAATTC | | | |
| ATRXcDNAF3 | CAGAGCCAGTGCTGAATGAA | 627 | | LMS35T |
| ATRXcDNAR3 | CAGAGCCAGAACAGGAATCA | | | |
| ATRXcDNAF4 | CAGTTGTTGCAGCAAAATAAGAA | 495 | | |
| ATRXcDNAR4 | TTCCAAAGCACAAGGTTTTTC | | | |
| ATRXcDNAF5 | TGGATGCTGTAAACAAAGAGAAA | 552 | | |
| ATRXcDNAR5 | CAGCACCTTTAATTGGGGAAT | | | |
| ATRXcDNAF6 | GGAGGTATTAAATCAAAAACTACAGC | 579 | LMS33T/LMS59T LMS63T/S832 | LMS59T/LMS63T |
| ATRXcDNAR6 | CAAAGTCTTATGGTTTGTATGAATTT | | | |
| ATRXcDNAF7 | TGAAATGCTAGCAATCCTCAAA | 773 | S825 | |
| ATRXcDNAR7 | CAGTTCCCTTTTTGCTCTGC | | | |
| ATRXcDNAF8 | AAGTACAGGATGGCTTATCTGATATT | 699 | | |
| ATRXcDNAR8 | GGGAGTTTCTCTTTTTCTCCTTG | | | |
| ATRXcDNAF9 | CAAAGTGGCTCATCATCATCTG | 684 | | |
| ATRXcDNAR9 | TTCACTGCTCACCTTTCTTCTG | | | |
| ATRXcDNAF10 | TGAGTGACGGAGAATCTGGA | 696 | | |
| ATRXcDNAR10 | CAGACTCACAGCAGCAATCC | | | |
| ATRXcDNAF11 | AAGATGCTTCACCCACCAAG | 802 | | LMS22T |
| ATRXcDNAR11 | TCTGCACACTGACCATTTTGA | | | |
| ATRXcDNAF12 | CATTGTATGGTTAATTTTATCAAGGAA | 761 | | LMS16T |
| ATRXcDNAR12 | TCAGCATCAGCATCTGTAACAA | | | |
| ATRXcDNAF13 | TTTAAAACTGGAAGAAAGTAAAGCTAC | 741 | | LMS48T/LMS60T |
| ATRXcDNAR13 | TCCCTCTTCTTCTTCTTTTCTGA | | | |
| ATRXcDNAF14 | TGGAGCGTCATTTTACTATGAAT | 559 | | |
| ATRXcDNAR14 | TCCTGGCTGGCTTGTCTACT | | | |
| ATRXcDNAF15 | ACAGTGTGACAGCAGTGAGGA | 545 | | |
| ATRXcDNAR15 | TTGAGTTCTGTTAAGTCATTGATTCC | | | |
| ATRXcDNALMS33F | GTGGACTTGGACAGGAAAACA | 690 | | LMS33T |
| ATRXcDNALMS33R | TGCTGTGTTTCTCATCTTCAGA | | | |
| ATRXcDNALMS35F | AACAATGTAGGTGGTGTGCG | 234 | | LMS35T |
| ATRXcDNALMS35R | TCTTATTTTGCTGCAACAACTGT | | | |
| ATRXBPcDNALMS66F1 | TCATCCTCTAGTTTGAAGCAAGG | 233 | | LMS66T |
| ATRXBPcDNALMS66R2 | CCTCAGCAAACTAACACAGG | | | |

**Table S6:** Primers used for *ATRX* cDNA screening and validation of mutations by Sanger sequencing related to methods.

*ATRX* forward and reverse primers used in cohort 2 for *ATRX* cDNA screening in both cohorts for some exon 9 *ATRX* DNA mutations and in cohort 1 for validation of cDNA mutations are presented. TD60°C PCR program described in Supplementary Table 5 legend was used. Primers used for cDNA validation of fusion transcript in LMS66 are also presented (primers in red). Tumors of cohort 1 in which each primer couple was used are indicated. For chohort 2 primers numbered from 1 to 15 were all used in all tumors to performed the ATRX RNA screen. bp: base pair, F: forward, R: reverse.

| Primer name | Primer sequence 5'-->3' | Tumor |
|---|---|---|
| ATRXex1F | CAGTGCATTTCTATCGTAACCG | S823 |
| DDX11L9ex1R | CTCTAGGCATGGCTCCTCTC | |
| ATRXex15F | TGAGAACAGAAACACAAAATGCT | S918 |
| RP1-279N11.1 75878708F | CACAACTGGCTCAACTGCTT | |
| ATRXex15F | TGAGAACAGAAACACAAAATGCT | |
| RP1-279N11.1 75879066F | TCATGTCAGTCTTTGTTGAGCA | |
| ATRXex27F | GTCCCTCATATCTCTGGACTTGA | |
| RP1-279N11.1 75913486F | TCTCTGTCCAGGCTTTCTTGA | |
| ATRXex27F | GTCCCTCATATCTCTGGACTTGA | M979 |
| FTXintronF | TGGAAGAATGTAAGGCCAGG | |
| ATRXex30F | CAGGTGGAGCGTCATTTTACTA | S831 |
| RNU6-974P81233619R | CAAGAATGTGGAGAGAGCCTTC | |
| ATRXex30F | CAGGTGGAGCGTCATTTTACTA | |
| RNU6-974P81166026R | GCAGTTGACACTTGCTACAATG | |
| DDX11L9ex1F | CTCTTAGCCCAGACTTCCCG | S823 |
| ATRXex1R | CAGAGTTACTTCCAGAACCACT | |
| RP1-279N11.1 75913486R | AGTGCTCTCTCTTGTTTATGGGA | S918 |
| ATRXex27R | CTCGATTAGCAGCTACCAGAT | |
| RP1-279N11.1 75878708R | TCAACTTTGCTGAGCCAAACT | |
| ATRXex15R | TCTTACCAAGGCCCATACAGT | |
| RP1-279N11.1 75879066R | GCCCTGGAGCTTACAATCAG | |
| ATRXex15R | TCTTACCAAGGCCCATACAGT | |
| FTXintronR | CAGGGAGCTTTCACATCAAACA | M979 |
| ATRXex27R | CTCGATTAGCAGCTACCAGAT | |
| RNU6-974P81233619F | TCACTGAGAACATTCAACCCC | S831 |
| ATRXex30R | GTTGATTACTCATTGCTGACAGG | |
| RNU6-974P81166026F | GAGGAAGACAAGTTTGGGCA | |
| ATRXex30R | GTTGATTACTCATTGCTGACAGG | |

**Table S7:** Combination of primers used to detect fusion transcript in cohort 2 related to methods.

First part of table indicates primer combination used to detect each fusion transcript. Second part describes primer combination used to detect potential reciprocal fusion transcript. TD60°C PCR program described in Supplementary Table 5 legend was used. F: forward, R: reverse, Ex: exon.

**Supplementary methods:**

***DNA extraction***

For both cohorts, genomic DNA from frozen samples was isolated using standard phenol-chloroform extraction protocol. DNA was quantified using Nanodrop 1000 spectrophotometer (Thermo Fisher Scientific, Waltham, MA, USA). Blood material was also available for cohort 1. Genomic DNA from blood samples was extracted using customized automated purification of DNA from compromised blood samples on the Autopure LS device according to the manufacturer's recommendations (9001340, Qiagen, Hilden, Germany) with increased centrifugation of 10 min for DNA precipitation and DNA wash.

***Whole genome sequencing and analysis***

Whole genome sequencing was performed only on cohort 1. To construct short-insert paired-end libraries, a no-PCR protocol was used with the TruSeq™DNA Sample Preparation Kit v2 (FC-121-2001/FC-121-2002, Illumina Inc., San Diego, CA, USA) and the KAPA Library Preparation Kit (KK8235, Kapa Biosystems, Basel, Switzerland). Briefly, 2 µg of genomic DNA were sheared on a Covaris™ E220, size-selected and concentrated using AMPure XP beads (A63880, Agencourt, Beckman Coulter, Brea, CA, USA) in order to reach a fragment size of 220 – 480 bp. Fragmented DNA was end-repaired, adenylated and ligated to Illumina-specific indexed paired-end adapters.

DNA sequencing was performed in paired-end mode, 2x100 bp or 2x125 bp according to flowcell version, in five or three sequencing lanes of HiSeq2000 flowcell v3 or v4 (Illumina Inc., San Diego, CA, USA) to analyze tumor or normal/constitutive samples and to reach minimal yield of 145 or 85 Gb, respectively. Two tumor samples were sequenced in 20 lanes to reach a minimal yield of 560 Gb. Image analysis, base calling and quality scoring of the run were processed using the manufacturer's Real Time Analysis software (RTA 1.13.48) and followed by generation of FASTQ sequence files by CASAVA (Illumina Inc., San Diego, CA, USA).

DNA reads were trimmed of the 5' and 3' low quality bases (phred cut-off 20, max trim size 30 nt) and sequencing adapters were removed with Sickle2 [44] (https://github.com/najoshi/sickle) and SeqPrep3 (https://github.com/jstjohn/SeqPrep), respectively. Then, DNA-curated sequences were aligned using Bowtie v2.2.1.0 [45], with the very sensitive option, on the Human Genome version hg19. Thus, aligned reads were filtered out if their alignment score was less than 20 or if they were duplicated PCR reads, with SAMtools v0.1.19 [46] and PicardTools v1.118 [47] (http://broadinstitute.github.io/picard/), respectively. SNV were detected by SAMtools mpileup v0.1.19 [46], with a minimum of 20 as phred quality score (-Q 20), and by bcftools call -Am (SAMtools v0.2.0, http://samtools.github.io/bcftools/call-m.pdf).

***RNA extraction, sequencing and analysis***

For both cohorts, RNA extraction was performed using standard TRIzol (15596026, Thermo Fisher, Waltham, MA, USA) / chloroform extraction (32211-1L, Fisher Scientific, Hampton, NH, USA) followed by 100% ethanol precipitation and RNA purification using the RNeasy Mini Kit (74104, Qiagen, Hilden, Germany) with DNase treatment (79254, RNase-Free DNase Set, Qiagen, Hilden, Germany). Total RNA was quantified using a Nanodrop 1000 spectrophotometer (Thermo Scientific, Waltham, MA, USA) and qualified with an Agilent 2100 Bioanalyzer (Agilent, Santa Clara, CA, USA) using the Agilent RNA 6000 Nano Kit (5067-1511, Agilent, Santa Clara, CA, USA) according to the manufacturer's instructions.
To control the sequencing quality, ERCC RNA Spike-In Mix (4456740, Life technologies, Carlsbad, CA, USA) were added to each RNA sample as recommended by the manufacturer. Analysis of this quality control was performed as previously described [18].

Regarding cohort 1, libraries from total RNA were prepared using the TruSeq®Stranded Total RNA Gold Library Preparation Kit (RS-122-2301, Illumina Inc., San Diego, CA, USA) according to the manufacturer's protocol. Briefly, 0.5 µg of total RNA was ribo-depleted using the Ribo-Zero Gold Kit. RNA fragmentation resulted in fragments of 80 – 450 nt, with a major peak at 160 nt. First-strand cDNA synthesis by random hexamers and reverse transcriptase was followed by second-strand cDNA synthesis, performed in presence of dUTP instead of dTTP. Blunt-ended double-stranded cDNA was 3´adenylated and Illumina-indexed adapters were ligated. Resulting libraries were enriched with 15 PCR cycles.

Construction of cohort 2 libraries was the same as previously described for frozen samples [18].

Libraries, of these two cohorts were sequenced on HiSeq2000 (Illumina Inc., San Diego, CA, USA) in paired-end mode with a read length of 2x75 bp using TruSeq SBS Kit v3-HS (FC-401-3001, Illumina Inc., San Diego, CA, USA). Image analysis, base calling and base quality scoring of the run were processed by integrated primary Real Time Analysis (RTA 1.13.48) software and followed by generation of FASTQ sequence files by CASAVA (v1.8, Illumina

Inc., San Diego, CA, USA). Library construction and RNA sequencing were performed at the Centro Nacional de Análisis Genómico (CNAG, Barcelona, Spain).

RNA bioinformatic analysis (alignment and expression quantification) for these two cohorts was performed as previously described [18]. Fusion transcripts were detected with Defuse v0.6.1 [48] as previously described [49].

Regarding cohort 1, SNV (Single Nucleotide Variant) were detected using samtools mpileup (SAMtools v0.1.19 [46]), with a minimum of 20 as phred quality score (-Q 20), and bcftools view -cvg (SAMtools v0.1.19 [50]). Regarding cohort 2, SNV were detected using samtools mpileup [46], with a minimum of 20 as phred quality score (-Q 20), and bcftools view -cgN [50]. Detected variants with fewer than 5 coverage reads were filtered out.

***Annotation of variants***

Regarding cohort 1, variants detected in constitutional, tumor DNA and tumor RNA were merged in the same file. Then, somatic variants were extracted with: (i) a minimum coverage of 14 reads in the tumor and 8 in the normal and (ii) a minimal allelic fraction of 0.3 in tumor and 0 in normal.

Since tumor and constitutional DNA were not available for cohort 2, candidate variants could not be filtered based on genotype differences between constitutional DNA and DNA/RNA tumor. Instead, filters applied were the same as for cohort 1 above, but with a minimum coverage of 5 reads due to a lower sequencing depth.

Variants were annotated using the Annovar v20160314 tool [51]. Variants were selected whose alternative allele frequency (AF) in the Caucasian population (CEU) is lower than 0.1%, as reported in the 1000Genome database [52]. Finally, variants were kept if they were localized in coding regions and were non-synonymous.

***Break Point Detection***

Structural variants (SV) were detected from paired tumor/normal whole genome high-quality sequencing data. Paired-end reads were aligned using Bowtie v2.2.1.0 [45], very sensitive local option allowing soft-clipped sequences. The algorithm has three main steps: i) identification of potential breakpoints, ii) characterization of the second side of the breakpoints, and iii) selection of high-confidence breakpoints. All parameters were set to analyze 60X tumor and 30X normal sequencing depth. Very conservative filters were used to minimize false positive detection.

i) Identification: at this step, reads with at least one soft-clipped end were analyzed as singletons. A position was considered as a potential breakpoint if it was covered by at least 4 soft-clipped reads, 5 soft-clipped bases (with at least two occurrences of two different bases), and if they represented more than 5% of the total amount of reads at this position in the tumor sample. We selected potential somatic events by discarding positions covered by at least 1 read and 1 base in a surrounding 5-nucleotide window in the normal sample. We refer to them as the "first side" of the breakpoint.

ii) Characterization: to determine the genomic positions of the soft-clipped sequence from selected reads, we used the UCSC blat server [53]. If no match was returned, the reverse complement sequence was pulled to test. If there was still no match, the BAM file was investigated for some soft-clip somatic position around the discordant or oversized-insert read mate (hereafter named abnormal) location from the first side of the breakpoint. Because of the small size of the soft-clipped sequence, multiple matches can be found. We used soft-clipped abnormal read mates to select matches with the most coherent chromosomic locations. We refer to them as the "second side" of the breakpoint.

iii) Selection: Positions detected from both the first and second sides (in a 5-nucleotide window) were defined as the common pool. We considered as artifacts (due to repeat regions for instance) couples of positions covered with reads and associated soft-clipped sequences separated by fewer than 15 nucleotides and discarded them. We classified the breakpoints in three groups: high-confidence breakpoints, breakpoints needing investigation, and unique position breakpoints. If a breakpoint was covered by reads and associated soft-clipped sequences having both positions belonging to the common pool, it was classified in the first group. If a breakpoint was covered by reads and associated soft-clipped sequences having only one of the positions belonging to the common pool, it was classified in the second group. Then the missing position was searched among the filtered positions. If it was present in the normal sample, the position was discarded and the breakpoint was completed otherwise. Finally, the third group corresponds to breakpoints with both sides outside the common pool and considered as unique: these were discarded. The sides of breakpoints were sorted according to their chromosomic positions to avoid duplicates.

For all the 67 LMS (cohort 1), the whole *ATRX* sequence obtained by WG sequencing was visualized using the Integrative Genomics Viewer (IGV, v2.6.3) [54] and soft-clipped reads were detected for 5 cases. *ATRX* fusion partner sequences and breakpoints location could be determined thanks to the blat function on the UCSC website.
